# Cell State Chaos Underpins the Evolution of SMARCA4-Deficient Dedifferentiated Endometrial Cancer

**DOI:** 10.64898/2025.12.01.690470

**Authors:** Mackenzie Brandon-Coatham, Julia Vassalakis, Hannah Plummer, Jasleen Kaur, Farzaneh Afzali, Zhihua Xu, Guihua Zhang, Jiahui Liu, Wei Wang, Einav Wajsbrot Renert, Tyler T. Cooper, Alan Dimitriev, Cierra Perron, Andrew Garven, Simatsidk Haregu Abebe, David M. Berman, Amber L. Simpson, Gane Wong, Gilles Lajoie, Felix K.F. Kommos, Andreas von Diemlin, Sheela Abraham, David J.H.F. Knapp, Alan Underhill, Martin Koebel, Cheng Han-Lee, Lynne-Marie Postovit

**Affiliations:** University of Alberta, Department of Oncology, Edmonton, AB, Canada; Queen’s University, Department of Biomedical and Molecular Sciences, Kingston, ON, Canada; Queen’s University, Sinclair Cancer Research Institute, Division of Cancer Biology and Genetics, Kingston, ON, Canada; Kingston Health Sciences Centre and Queen’s University, Department of Pathology and Molecular Medicine, Kingston, ON, Canada; Université de Montréal, Centre de recherche du Centre hospitalier de l’Université de Montréal, Department of Obstetrics and Gynecology, Montréal, QC, Canada; University of Alberta, Department of Biological Sciences, Edmonton, AB, Canada; Queen’s University, School of Computing, Kingston, ON, Canada; University of Western Ontario, Department of Biochemistry, London, ON, Canada; University Hospital of Heidelberg, Institute of Pathology, Heidelberg, Germany; University Hospital of Heidelberg, Department of Neuropathology, Heidelberg, Germany; Université de Montréal, Department of Pathology and Cell Biology, Montréal, QC, Canada; University of Calgary, Department of Pathology and Laboratory Medicine, Foothills Medical Centre, Calgary, AB, Canada; University of Alberta, Department of Laboratory Medicine and Pathology, Edmonton, AB, Canada

## Abstract

Dedifferentiated endometrial carcinoma (DDEC) is a histologically unique cancer type, wherein well-differentiated regions lie adjacent to morphologically distinct, high-grade lesions that are histologically undifferentiated. Previous studies have determined that in nearly half of the cases dedifferentiation is associated with the genomic inactivation of *SMARCA4*, a catalytic subunit belonging to the SWI/SNF chromatin remodelling complex (SWI/SNF CRC), suggesting that SMARCA4 loss causes dedifferentiation. Herein, using gene editing, we reveal that when serially passaged in mice, SMARCA4-deficient endometrial cancer cells repeatably and predictably generate heterogeneous admixtures of differentiated and undifferentiated cells, resembling human DDEC. Surprisingly, despite this metamorphosis, SMARCA4 loss does not induce lineage plasticity nor reprogramming to a less differentiated fate. Rather, single-cell sequencing combined with barcoding demonstrated that *SMARCA4* loss induces a dysregulated epigenome that allows cells to randomly move through cellular states that are otherwise shared with *SMARCA4-*expressing well-differentiated cancer cells. This finding was validated using a cohort of patient samples, such that epithelial fate markers (E-CADHERIN) can be detected in morphologically undifferentiated cells. Collectively, this work constitutes the first repeatable model of human dedifferentiated cancer and suggests that histological dedifferentiation is not due to the acquisition of a stem cell-like fate. Rather, undifferentiated tissue emerges due to epigenomic dysfunction concomitant with the arbitrary movement of cancer cells between cellular states.

## INTRODUCTION

Neoplastic progression is accompanied by the acquisition of cellular plasticity, wherein cancer cells manifest a continuum of lineage-specific phenotypes^1^. Cancer cell plasticity leads to heterogeneous and dynamic tumours that are better able to evade the immune system, survive therapy, and spread^2^. Mechanistically, plasticity occurs in response to microenvironmental cues, such as hypoxia, which triggers coordinated reprogramming of the epigenome, transcriptome, and translatome^3^. While cellular plasticity is thought to be acquired cumulatively during progression, there are histopathologies wherein the tumour can undergo abrupt metamorphosis, with well-differentiated regions adjacent to morphologically distinct, high-grade lesions that are histologically undifferentiated. These cancers, including dedifferentiated chondrosarcoma^4^, dedifferentiated liposarcoma^5,6^, metaplastic breast cancer^7–9^, rhabdoid lung carcinoma^10^, dedifferentiated endometrial carcinoma (DDEC) and dedifferentiated ovarian carcinoma^11–13^ exemplify an extreme state of cellular plasticity and can thus be used to understand the mechanisms by which this phenomenon is acquired and sustained. It is generally believed that dedifferentiation is a clonal event that occurs due to the acquisition of genetic abnormalities^14–17;^ however, the dynamic interplay between mutations and microenvironmental mediators of cellular plasticity has not yet been described.

DDEC, characterized typically by the presence of a well-differentiated endometrial carcinoma adjacent to an undifferentiated carcinoma^18^, is associated with poor outcomes due to therapy resistance and metastatic spread^19^. Genomic studies comparing undifferentiated regions with their well-differentiated precursor lesions revealed that approximately two-thirds of dedifferentiation events occur concomitantly with deficiencies in the SWItch/Sucrose-Non-Fermentable (SWI/SNF) chromatin remodelling complex; most notably, the loss of SMARCA4 or the concurrent loss of ARID1A and ARID1B^20^. The SWI/SNF complex modulates chromatin architecture and regulates gene expression by repositioning nucleosomes^21^. SMARCA4 is the ATPase subunit of one SWI/SNF complex, while ARID1A and ARID1B are essential DNA-binding subunits^22^. Therefore, mutations that impair the SWI/SNF complex, as observed in the undifferentiated areas of DDEC lesions, are expected to impact chromatin remodelling significantly. The loss of SMARCA4 has been shown to promote early metastatic events and to reduce the expression of lineage-specific markers in various cancers, including non-small cell lung carcinoma^23^. Although SMARCA4 loss occurs in nearly half of DDEC cases^20^, the extent to which it may drive dedifferentiation in endometrial carcinomas remains to be thoroughly investigated.

Herein, we demonstrate that endometrial cancer cells engineered to have *SMARCA4* knocked out form tumours in mice that recapitulate human DDEC. In contrast to wild-type counterparts, these SMARCA4-deficient tumours contain differentiated carcinoma juxtaposed to undifferentiated carcinoma and are resistant to chemotherapy. Using single-cell multiome sequencing together with barcoding^24^, we determined that wild-type and knockout cells exist in similar states. However, while wild-type cells appear to have ordered trajectories and express each cluster proportionally during passaging, knockout cells move erratically between cellular states. This suggests that the appearance of dedifferentiated tissue may be due to the arbitrary and transient acquisition of different states (cell state chaos) rather than dedifferentiation per se. This was confirmed using a cohort of DDEC patient tissues, wherein E-CADHERIN (*CDH1*), a marker of differentiated endometrial cancer^25^, was randomly expressed in cells throughout histologically undifferentiated regions. Collectively, these results suggest that the loss of SMARCA4 did not induce clonal dedifferentiation. Hence, the undifferentiated lesions present in DDECs do not occur due to lineage plasticity. Rather, the tumour appears undifferentiated as a consequence of cell state chaos, wherein epigenetically dysregulated cells arbitrarily move between cellular phenotypes.

## RESULTS

### *SMARCA4* loss initially induces a senescent-like phenotype in mismatch repair (MMR)-deficient endometrial cancer cells

To investigate whether the loss of SMARCA4 can recapitulate the aggressive phenotype found in the undifferentiated region of DDEC patients, we used CRISPR-Cas9 to genocopy *SMARCA4* loss-of-function mutations in HEC116 and HEC59 human endometrial cancer cell lines. These cell lines are classified as microsatellite instability (MSI)/hypermutated, which is the most prevalent molecular context for DDEC and undifferentiated endometrial carcinoma (UDEC) cases^26^, particularly among SWI/SNF-deficient DDEC^27^. Two different CRISPR-Cas9 methodologies were applied to generate the SMARCA4 knockout (KO) cells: a plasmid-derived system (PD) and a direct delivery ribonucleoprotein complex system (RNP) (**Fig. 1a**). The presence of indel mutations confirmed the deletion of *SMARCA4* (**Supplementary Fig. 1a**) and the loss of protein expression in the cell lines was validated by immunohistochemistry (IHC) (**Fig. 1a**). The loss of SMARCA4 induced a significant shift in the transcriptome of HEC116 SMARCA4 KO cells compared to their WT counterparts (**Supplementary Fig. 1b,c**), causing the upregulation and downregulation of 3,065 and 1,607 transcripts, respectively (**Fig. 1b**). Enrichment analyses showed pathways related to estrogen response and adherent junctions of epithelial cells to be downregulated after the loss of SMARCA4. In contrast, pathways related to epithelial-to-mesenchymal transition, response to hypoxia, stemness, and extracellular matrix modulation were upregulated (**Supplementary Fig. 1d**). By qPCR, *CDH1*, *CLDN3* and *ER* gene expression were downregulated, and *OCT4* was upregulated in *SMARCA4* KO cells, corroborating some bulk RNA-sequencing findings (**Supplementary Fig. 1e**). Mass Spectrometry (MS)-based proteomics revealed a strong positive correlation between transcripts and proteins found up- or down-regulated in HEC116 SMARCA4 KO cells (r = 0.85; 95% CI 0.81-0.87) (**Supplementary Fig. 1f**). Proteome analysis on cell lysates revealed that 459 and 361 proteins were upregulated and downregulated, respectively, after SMARCA4 loss in HEC116 cells (**Fig. 1c**). Functionally, these proteins were associated with processes such as metabolism, extracellular matrix remodelling, and adaptive stress responses (**Fig. 1c; Supplementary Fig. 2**).

**Figure 1.**
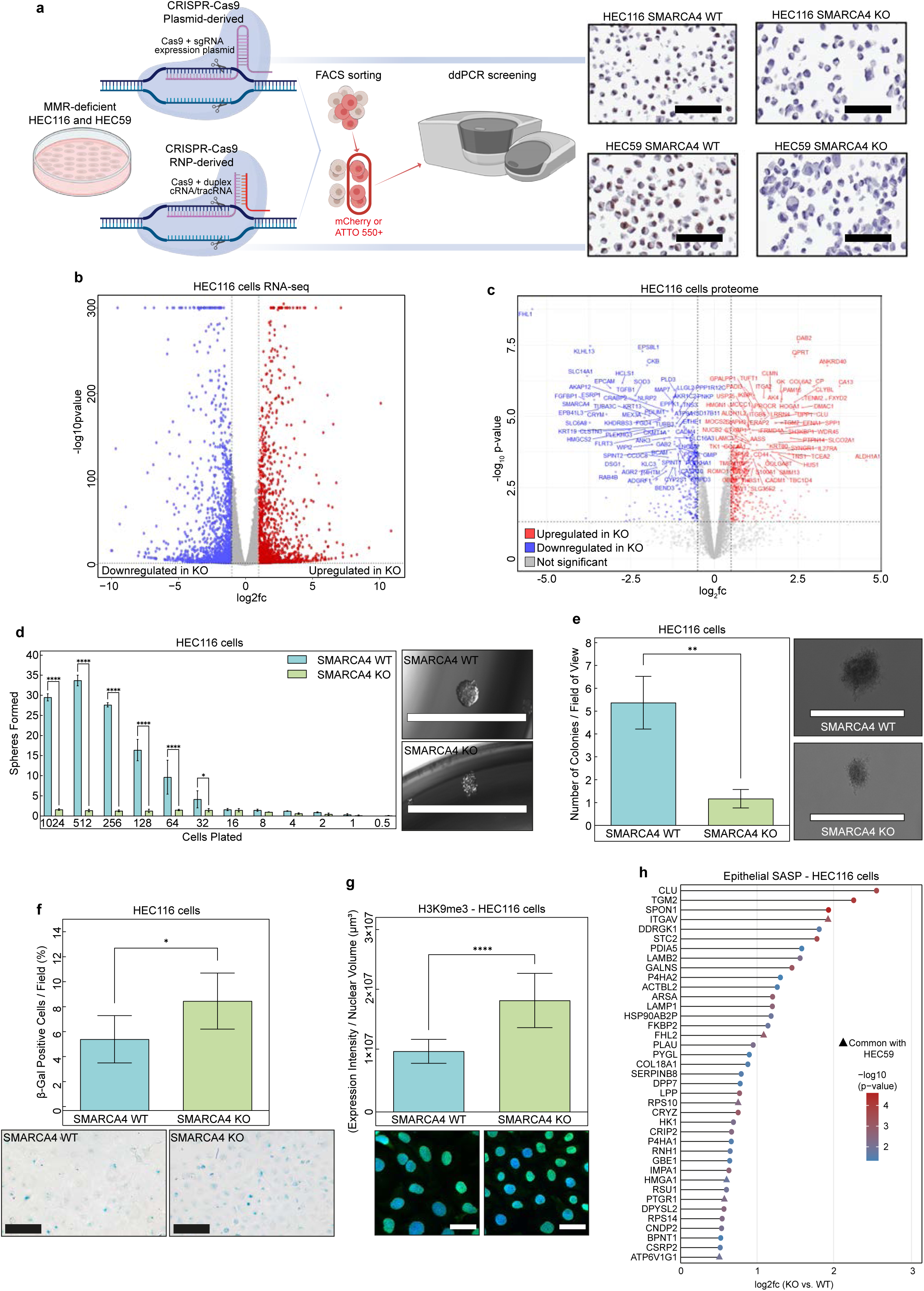
SMARCA4 loss initially induces a senescent-like phenotype in endometrial carcinoma cell lines. (a) CRISPR-Cas9 workflow to generate SMARCA4-deficient human endometrial cancer cell lines. HEC116 and HEC59 cells were transfected with a Cas9+gRNA plasmid or a Cas9 cRNA/tracRNA RNP complex, respectively. Edited mCherry⁺ or ATTO550⁺ cells were enriched by flow cytometry, expanded as single-cell clones, and screened by droplet digital PCR (ddPCR). Clones were validated by Sanger sequencing, and SMARCA4 immunostaining confirmed protein loss. Scale bar: 100 μm. (b) Volcano plot of differential gene expression (bulk RNA-seq). Upregulated genes in HEC116 SMARCA4 KO versus WT are shown in red, downregulated in blue, and non-significant in grey (*p* < 0.05, |log₂FC| > 1). (c) Volcano plot of MS-based whole-cell proteome. Upregulated proteins in HEC116 SMARCA4 KO compared to WT are shown in red, downregulated in blue, and non-significant in grey (*p* < 0.05, |log₂FC| > 0.5), with gene names indicated for significant proteins. (d) Limiting dilution sphere formation assay. Single-cell suspensions of HEC116 WT and SMARCA4 KO cells were seeded at decreasing densities in ultra low-attachment (ULA) plates and cultured in minimal media for up to two weeks. Data represent eight technical replicates from three independent experiments. Multiple t-tests with Holm-Šidák correction assessed statistical significance (\**p* < 0.05, \*\*\*\**p* < 0.0001). Representative images are shown (right). Scale bar: 1000 μm. (e) Anchorage-independent growth assay. HEC116 WT and SMARCA4 KO cells were embedded in 0.7% agarose-containing media and cultured for up to two weeks. Colonies were quantified from ten fields of view across three independent experiments. Statistical significance was assessed by unpaired t-test (\*\**p* < 0.01). Representative images are shown (right). Scale bar: 200 µm. (f) Senescence-associated β-galactosidase (SA-β-gal) assay. The percentage of SA-β-gal–positive HEC116 WT and SMARCA4 KO cells was quantified from six randomly selected fields. Statistical significance was assessed by unpaired t-test (\**p* < 0.05). Representative images are shown (bottom). Scale bar: 50 µm. (g) Immunofluorescence quantification of H3K9me3. HEC116 WT and SMARCA4 KO cells were analyzed for total H3K9me3 signal intensity per cell, normalized by cell volume, across two fields of view. Representative images are shown (bottom). Statistical significance was assessed by unpaired t-test (\*\*\*\**p* < 0.0001). Scale bar: 10 µm. (h) MS-based secretome analysis. Proteins in CM of HEC116 SMARCA4 KO compared to WT with log₂FC > 0.5 and *p* < 0.05 were compared to the epithelial SASP signature (http://saspatlas.com). Colour scale indicates p-values of upregulated proteins. Proteins upregulated in both HEC116 and HEC59 SMARCA4 KO vs. WT are marked with ▴.

Tumoursphere formation and anchorage-independent growth are surrogates for cancer cell self-renewal and tumourigenic potential. Since SMARCA4 loss is associated with dedifferentiated regions of DDECs, we reasoned that deleting SMARCA4 would enhance these tumourigenic phenotypes. However, in comparison to WT cells, HEC116 and HEC59 SMARCA4 KO cells formed fewer and smaller spheres (**Fig. 1d; Supplementary Fig. 3a**) and were less able to grow on soft agar (**Fig. 1e; Supplementary Fig. 3b**). Cancer-driving events such as the acquisition of RAS activating mutations^28^ or the loss of ARID1A can initially induce senescence concomitant with cell cycle arrest and senescence-associated secretory phenotype (SASP)^29^. Accordingly, β-galactosidase assays showed higher enzyme activity in all SMARCA4 KO cells (**Fig. 1f; Supplementary Fig. 3c**), indicating a senescent-like state, and p21 (a marker of cell cycle arrest) accumulated in HEC59 SMARCA4 KO cells (**Supplementary Fig. 3d**). As compared to WT cells, SMARCA4 KO HEC116 cells also had an increase in senescence-associated heterochromatin foci (SAHFs) that were enriched for the repressive mark H3K9me3^30^ (**Fig. 1g; Supplementary Fig. 3d**). Using MS-based proteomics of conditioned media (CM), we determined that, compared to WT cells, SMARCA KO HEC116 cells secreted 39 proteins (e.g., CLU, TGM2, and SPON1) (**Fig. 1h**) that are associated with the SASP in epithelial cells (http://saspatlas.com)^31^. In comparison to WT counterparts, SMARCA KO HEC59 cells exhibited a similar increase in the secretion of 56 proteins associated with the SASP (**Supplementary Fig. 3e**). Collectively, these results indicate that the loss of SMARCA4 causes significant transcriptional reprogramming in endometrial carcinoma cell lines and that this results in an altered proteome. However, this reprogramming does not induce the aggressive phenotypes observed in patients diagnosed with DDEC. The absence of the expected pro-tumourigenic phenotypes is likely due to the acquisition of a senescence-like state in endometrial carcinoma cell lines following SMARCA4 loss.

### Following *in vivo* serial passaging, SMARCA4 loss in endometrial cancer cells enables the evolution of poorly differentiated lesions that recapitulate clinical DDEC

Given the association of SMARCA4 loss and DDEC in patients, we hypothesized that the initial senescence-like phenotype induced by SMARCA4 loss could be overcome upon exposure to the tumour microenvironment. Accordingly, we injected HEC116 and HEC59 WT and SMARCA4 KO cells into NOD.Cg-Prkdc*^scid^*Il2rg*^tm1Wjl^*/SzJ (NSG) mice, and then serially passaged tumour fragments into recipient mice for at least four generations (**Fig. 2a**). Corroborating our *in vitro* tumoursphere findings, SMARCA4 KO initial generation (F0) tumours were smaller as compared to WT. In HEC116 CDX, measurements on day 104 post-injection showed median tumour volumes of 161 mm³ for SMARCA4 KO versus 544 mm³ for WT, with KO tumours growing more slowly (F0 tumour growth rate (TGR): KO = 0.070 ± 0.017 vs. WT = 0.113 ± 0.015) (**Fig. 2a; Supplementary Fig. 4a**). In HEC59, tumour growth was significantly delayed, with WT tumours reaching a median volume of 479 mm³ after 36 days and SMARCA4 KO tumours reaching 280 mm³ at 151 days post-injection (**Supplementary Fig. 4b**). In the first passage (F1), HEC59 KO tumours maintained a slower growth rate relative to WT tumours (F1 TGR: KO = 0.06 ± 0.01 vs. WT = 0.13 ± 0.007) (**Supplementary Fig. 4c**). However, after subsequent passages, HEC116 and HEC59 SMARCA4 KO tumours grew at a rate equal to or faster than WT tumours, suggesting that the senescence-like phenotype emerging upon SMARCA4 loss can be overcome (**Supplementary Fig. 4a, c**). Indeed, after serial passaging, cells from SMARCA4 KO tumours efficiently formed spheroids with a grape-like morphology (**Fig. 2b**), exhibited lower β-galactosidase activity compared to SMARCA4 KO cells in culture or WT cells similarly passaged (**Fig. 2c**), and showed comparable H3K9me3 expression to WT cells (**Supplementary Fig. 4d**).

**Figure 2.**
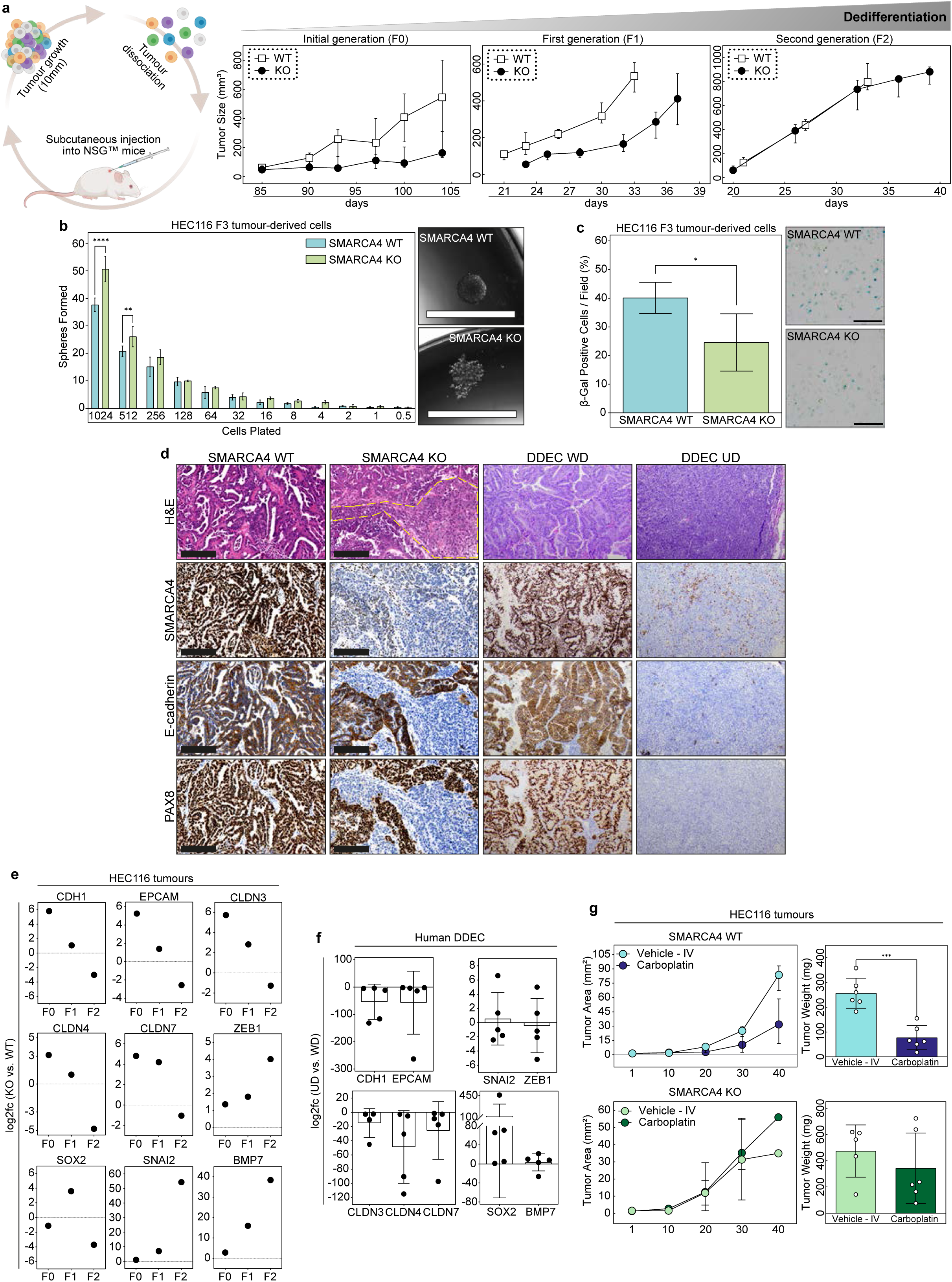
SMARCA4 loss recapitulates human DDEC upon in vivo serial passages. (a) Schematic representation of *in vivo* serial passaging of HEC cells. Left: tumour cells were injected subcutaneously into NSG mice, dissociated at ∼10 mm, and reimplanted into subsequent mice. Right: tumour growth kinetics of HEC116 WT and SMARCA4 KO cells across initial (F0), first (F1), and second (F2) passages (n = 6 mice per group). (b) Limiting dilution sphere formation assay of serially passaged cells. Single-cell suspensions of third-generation (F3) HEC116 WT and SMARCA4 KO CDX-derived cells were seeded at decreasing densities in ULA plates and cultured in minimal media for up to two weeks. Data represent eight technical replicates from three independent experiments. Statistical significance was assessed by multiple t-tests with Holm-Šidák correction (\*\**p* < 0.01, \*\*\*\**p* < 0.0001). Representative images are shown (right). Scale bar: 1000 µm. (c) SA-β-gal staining of serially passaged cells. The percentage of SA-β-gal–positive HEC116 WT and SMARCA4 KO F3 CDX-derived cells was quantified from five images across four independent experiments performed in duplicate. Statistical significance was assessed by unpaired t-test (\**p* < 0.05). Representative images are shown (right). Scale bar: 200 µm. (d) Histology of serially passaged tumours compared to human DDEC. Representative H&E and IHC images of HEC116 WT and SMARCA4 KO tumours are shown alongside human DDEC samples. IHC staining for SMARCA4, E-CADHERIN, and PAX8 highlights loss of SMARCA4 and reduced E-CADHERIN and PAX8 in undifferentiated (UD) regions of SMARCA4 KO and DDEC tumours, whereas WT tumours and well-differentiated (WD) regions of DDEC maintain expression. Scale bar: 200 µm. (e-f) Nanostring analysis of EMT and stemness-associated genes in serially passaged tumours and human DDEC. (e) Log₂FC of selected genes in HEC116 SMARCA4 KO versus WT tumours across F0, F1, and F2 passages. (f) Log₂FC of the same genes between paired UD and WD tumour regions in human DDEC. (g) Effect of carboplatin on tumour growth and weight. Tumour area over time (left) and tumour weight at endpoint (right) are shown for HEC116 WT and SMARCA4 KO tumours treated with carboplatin or vehicle (PBS). Tumour growth curves were compared using nonparametric Kolmogorov-Smirnov tests; tumour weights were compared by t-test (n = 6; \*\*\**p* < 0.001).

We next sought to determine whether SMARCA4 loss could induce histological dedifferentiation. Not uncommon for xenograft models, hematoxylin and eosin (H&E) staining showed that HEC59 tumours are poorly differentiated and do not have epithelial-like structures, making this a poor model for studying dedifferentiation (**Supplementary Fig. 4e**). In contrast, HEC116 WT cells formed tumours with endometrioid glandular-like features and this histology was maintained in all passages (**Fig. 2d; Supplementary Fig. 4f**). Astonishingly, despite being clonally derived and thus expected to yield homogeneously undifferentiated tumours, by the second passage SMARCA4 KO tumours began to recapitulate the histopathology of DDEC, with endometrioid regions juxtaposed to undifferentiated lesions (**Fig. 2d**). The extent of dedifferentiation appeared to increase with each passage. In clinical DDEC, the loss of PAX8 and E-CADHERIN marks dedifferentiated regions^32,33^. Accordingly, concomitant to SMARCA4 loss, the undifferentiated portions of SMARCA4 KO tumours showed reduced levels of E-CADHERIN and PAX8. In contrast, the more differentiated portion of SMARCA4 KO tumours retained expression of these proteins (**Fig. 2d**). This DDEC-like histology appeared in all SMARCA4 KO tumours, suggesting that it was not a random event (**Fig 2d; Supplementary Fig. 4g**). Using NanoString technology, we next analyzed the expression of mRNA targets related to EMT and stemness in HEC116 SMARCA4 WT versus KO CDX tumours throughout serial passaging (as the dedifferentiated component increased) and in the endometrioid and undifferentiated portions of human clinical DDEC samples. We found that, in comparison to WT tumours or the well-differentiated regions of human DDECs, SMARCA4 KO tumours and the undifferentiated regions of human DDEC were characterized by the loss of epithelial markers (e.g. *E-CADHERIN*, *EPCAM* and *CLDN3, 4* and *7*) (**Fig. 2e,f**). While transcription factors related to EMT (e.g. *ZEB1* and *SNAI2*) were upregulated in SMARCA4 KO versus WT tumours, the same was not observed in all the poorly versus well-differentiated components of human tissues (**Fig. 2e,f**). Certain morphogenic genes (e.g., *BMP7*) were increased in the undifferentiated portion of human DDECs, and in serial passages of HEC116 SMARCA4 KO tumours (**Fig. 2e,f**). However, *SOX2*, a gene associated with stem-like cells, was heterogeneously expressed across samples (**Fig. 2e,f**). Collectively, these results suggest that epithelial genes are downregulated in SMARCA4-deficient tumours upon dedifferentiation, but that certain genes (e.g. *SOX2*) may be regulated independently of dedifferentiation. Patients with SWI/SNF-deficient DDEC experience resistance to chemotherapy^18^. To determine whether this was recapitulated in the HEC116 model, mice harbouring SMARCA4 WT and KO F3 tumours were treated with 60mg/kg carboplatin or with vehicle. Mice were dosed every week for a total of five weeks, starting 14 days post-CDX implantation. While carboplatin reduced the final tumour weight and slowed tumour growth in mice harbouring WT tumours, it did not perturb the growth of SMARCA4 KO tumours (**Fig. 2g**). Hence, tumours derived from HEC116 SMARCA4 KO cells can faithfully recapitulate human DDEC. This represents the first repeatable model of endometrial cancer dedifferentiation.

### Histological undifferentiation of SMARCA4 KO tumours is not stochastic and is accompanied by sustained alterations in DNA methylation

We next wanted to determine the mechanisms governing dedifferentiation in SMARCA4 KO tumours. Clinically, dedifferentiation is posited to be clonal, emerging stochastically due to a mutation. To test the possibility that dedifferentiated SMARCA4 KO cells acquired specific mutations, we conducted whole-exome sequencing (WES) and then analyzed the tumour mutational burden (TMB) and mutational profiles in HEC116 WT and SMARCA4 KO cells prior to passaging in a mouse and after three sequential passages. Nonsynonymous TMB ranged from 11.65 to 12.15 mutations/megabase. The relatively high TMB, despite stringent filtering, is indicative of the HEC116 cell model which is MSI/hypermutated. The TMB was consistent between cells in culture versus cells derived from tumours (**Fig. 3a**). Mutations were identified in 575 genes across all samples, with 4.9-5.1% of mutated genes corresponding to cancer-related genes (COSMIC databases). Since the CRISPR-induced SMARCA4 mutation is detected in our analysis as germline (present in cells prior to passaging) rather than somatic (acquired or lost in tumours), we analyzed both germline and somatic nonsynonymous variants in cancer-related genes to enable comparisons of mutational profiles between WT and KO cells and across different microenvironments. While there were some differences in WT versus KO cells (likely due to clonal selection during CRISPR workflows), neither model appeared to acquire significant de novo variants during *in vivo* passaging (**Fig. 3b; Supplementary Fig. 5**). In the cases where a mutation was acquired, it was at a very low frequency that did not match the much higher levels of dedifferentiation observed in the tumours. Overall, mutational landscapes, including TMB and mutational profiles, were stable across samples, indicating minimal genomic adaptation during *in vivo* serial passaging. Given this finding, it is likely that dedifferentiation is occurring via epigenetic mechanisms.

**Figure 3.**
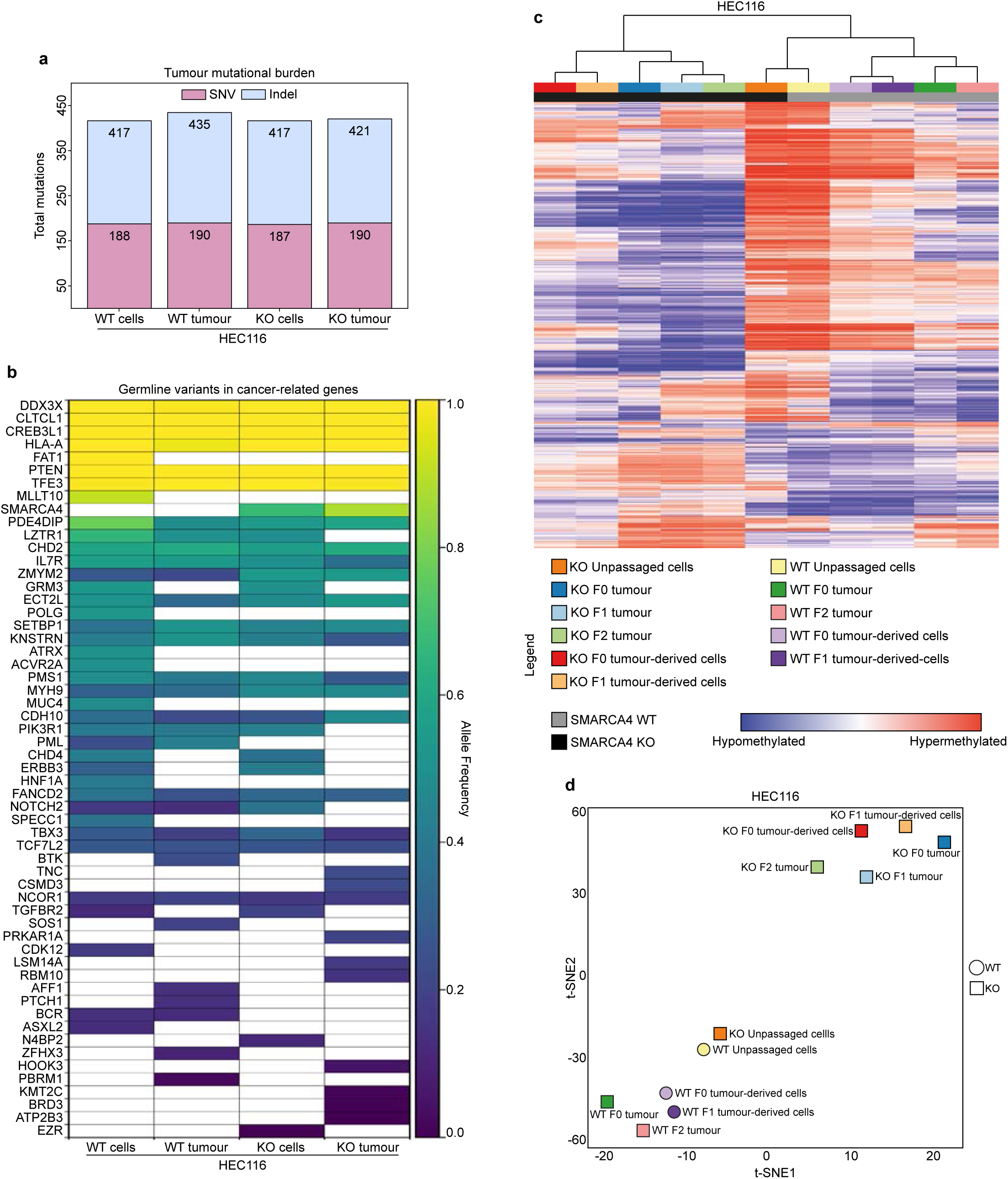
Epigenetic mechanisms mediate dedifferentiation in SMARCA4 KO tumours. (a) Tumour mutational burden (TMB) in HEC116 WT and SMARCA4 KO cells before and after three serial passages in mice, calculated from whole-exome sequencing (WES) data. Stacked bars show the total number of nonsynonymous mutations per sample, with single-nucleotide variants (SNVs) and insertions/deletions (indels) indicated within each bar. (b) Heatmap of allele frequencies of nonsynonymous variants in cancer-related genes in HEC116 WT and SMARCA4 KO cells, highlighting germline variants including the CRISPR-induced SMARCA4 mutation. Genes are defined according to the COSMIC Cancer Gene Census. Colours indicate allele frequency, with white representing absence of a mutation. (c) Global DNA methylation patterns in HEC116 WT and SMARCA4 KO samples. Hierarchical clustering of CpG methylation was performed across unpassaged cell lines, serially passaged tumours *in vivo*, and tumour-derived cells cultured *in vitro*. Columns represent individual samples and rows represent CpG sites. (d) t-distributed Stochastic Neighbor Embedding (t-SNE) analysis of genome-wide DNA methylation in HEC116 WT and SMARCA4 KO samples, including unpassaged cell lines, serially passaged tumours *in vivo*, and tumour-derived cells cultured *in vitro*.

Several studies have shown that SMARCA4 loss is associated with changes in global DNA methylation^34–38^. Accordingly, we next measured genomic DNA methylation patterns in HEC116 WT and SMARCA4 KO cells prior to passaging, in tumours, and in cells derived from tumours and then cultured *in vitro*. Using hierarchical and consensus clustering, we found that prior to *in vivo* passaging, there were few global differences between WT and SMARCA4 KO cells, and that these unpassaged cells clustered together (**Fig. 3c**). Tumour growth in mice caused hypomethylation in both SMARCA4 KO and WT cells; however, the patterns of methylation differed between the samples. Notably, methylation patterns of HEC116 WT cells remained similar between serial passages in a mouse and were largely unaffected by culture *in vitro* versus *in vivo*. However, SMARCA4 KO cells moved farther away from the original KO cells in culture with each additional passage and did not return to this state upon recovery *in vitro* (**Fig. 3d**). Collectively, these results suggest that the dedifferentiation observed in SMARCA4 KO does not occur stochastically and is associated with dysregulation of the epigenome that is precipitated by *in vivo* tumour formation.

### SMARCA4 KO tumour cells do not undergo unidirectional dedifferentiation

To establish the trajectory by which SMARCA4 KO cells become undifferentiated, we next employed multiome scRNA-seq and scATAC-seq in cells transduced with a barcode library prior to serial passaging, to ascertain how each clone evolved. Given the large epigenomic differences observed between cells in culture and those from tumours (**Fig. 3**), we elected to focus on cells extracted from the initial tumours (F0) and from subsequent serial passages (F1, F2). To validate that dedifferentiation does not occur due to alterations in mutational frequencies or to the outgrowth of a specific clone, we measured barcode frequencies over time. We determined that WT and KO tumours contained a similar number of clones and that these were maintained throughout the passages (**Fig. 4a**). As an orthogonal approach, we examined mitochondrial evolution and determined that WT and SMARCA4 KO tumours exhibited a similar level of entropy (**Fig. 4b**). Collectively, these results further support the hypothesis that dedifferentiation does not occur due to clonal selection following a stochastic mutational event. Unsupervised clustering of scRNA-seq data using Harmony^39^ for batch correction showed that except for one small cluster (C5), SMARCA4 KO and WT HEC116 cells exist in similar transcriptional states (**Fig. 4c; Supplementary Fig. 6**). While the clusters (particularly C5) had different transcription factor activities (**Fig. 4d**), the only statistically significant pathway that emerged was related to hypoxia (**Fig. 4e**). Cells expressing a hypoxic signature were present in both KO and WT cells, and likely represent cells isolated from hypoxic regions of the tumour. C5 was present only in SMARCA4 KO tumours and could be characterized as *CDH1* and *PAX8* negative; *BMP4* and *DLX1* positive (**Fig. 4f**). To validate these results, we employed three additional batch correction techniques (CCA^40,41^, RPCA^41^ and MNN^42^) (**Supplementary Fig. 7a**). Similar results were obtained with all methods, except for CCA, which did not isolate C5 as a separate cluster (**Supplementary Fig. 7a,b**). Rather, CCA incorporated the SMARCA4-KO-specific cells (CDH1^-^, PAX8^-^; BMP4^+^, DLX1^+^) into clusters shared with WT cells (**Supplementary Fig. 7c**), suggesting that these cells may not represent a distinct lineage. Results from an integrated scATAC-seq analysis demonstrated that in WT tumours, all transcriptional states occurred within a relatively homogeneous state of chromatin occupancy (**Fig. 4g**). In contrast, greater heterogeneity was observed in SMARCA4 KO tumours, with numerous chromatin states contributing to each transcriptional cluster (**Fig. 4g**). This suggests that in WT tumours, the transcriptional states were driven by changes in signaling rather than lineage plasticity and that in KO cells, the cells were able to manifest similar transcriptional patterns, despite having an altered epigenomic landscape. The C5 transcriptional cluster was characterized by a distinct chromatin occupancy. However, gene set enrichment analysis of this KO-specific cluster suggests that it is not enriched for stem cell factors nor for any specific lineage (**Supplementary Fig. 8**). Moreover, in contrast to the proportion of histological dedifferentiation, which increased with each sequential passage of SMARCA4 KO tumours, the size of cluster C5 did not increase over time and emerged in F0 tumours. Thus, we do not believe that it is a cancer stem cell population or that it represents the dedifferentiated component of DDEC lesions. Barcode analysis demonstrated that each clone had the ability to exist in all clusters (**Fig. 4h; Supplementary Fig. 9**). However, while WT clones maintained similar ratios of each cluster throughout passaging, KO clones were more heterogeneous. This was confirmed using lineage trajectory analysis (scVelo)^43^, which showed that WT clones moved directionally between states and rested mostly in an end state (**Fig. 4i**). In contrast, KO clones appeared to move randomly, with few cells existing at a clear end state. This analysis further confirmed that the C5 cluster did not act as a progenitor population and did not represent the dedifferentiated state, as it emerged as a late event and did not seem to occur due to unidirectional reprogramming from the other cellular states. Collectively, these results suggest that SMARCA4 KO does not promote directional dedifferentiation. Rather, it appears to enable epigenomic chaos, leading to free movement between cellular states.

**Figure 4.**
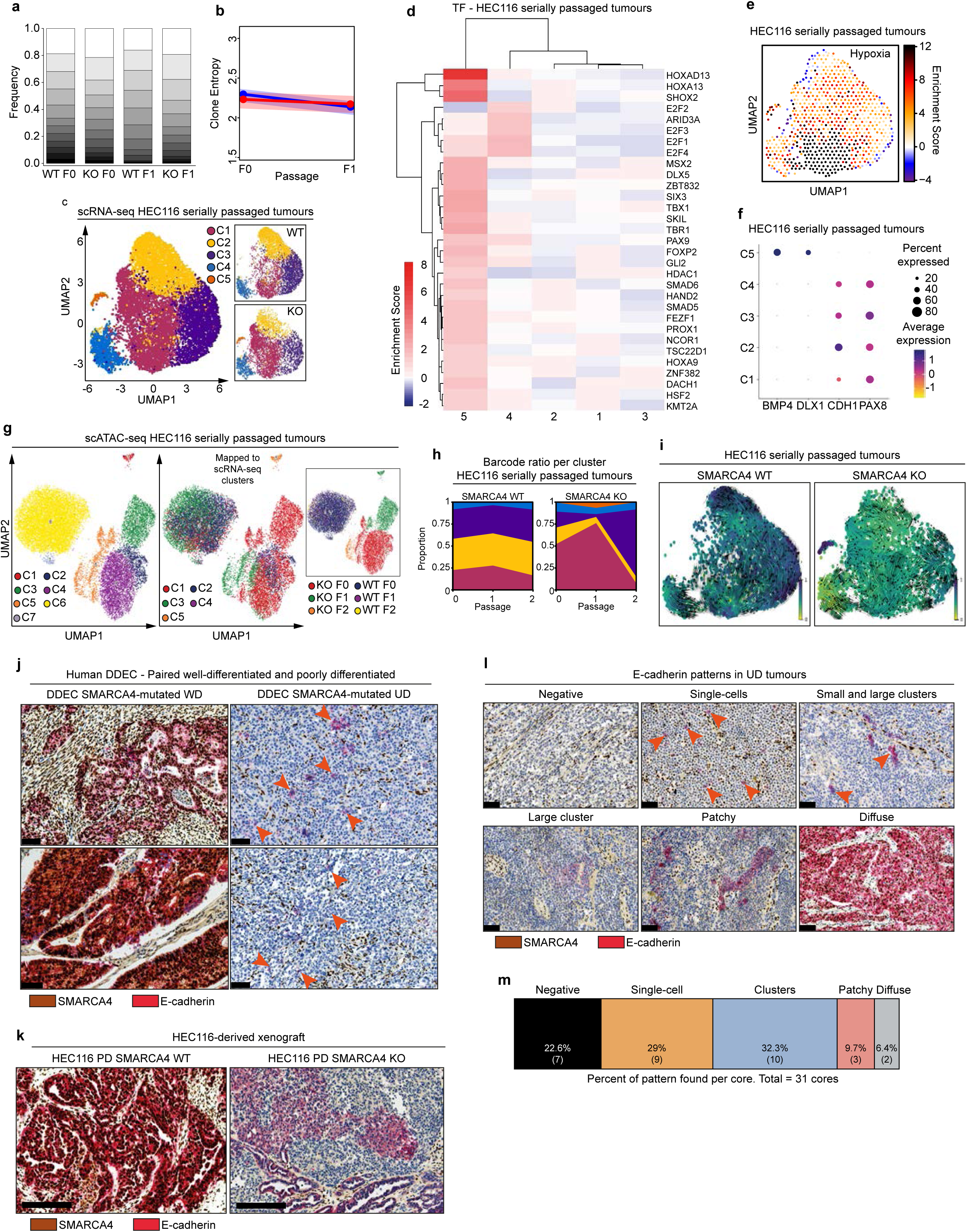
SMARCA4 deficiency induces cell state chaos rather than directional reprogramming. (a) Relative frequencies of lentiviral barcodes in serially passaged tumours. The proportions of individual barcodes are shown for HEC116 WT and SMARCA4 KO F0 and F1 tumours, illustrating that clonal composition stays consistent during serial passaging. (b) Clonal diversity measured by mitochondrial variant entropy in serially passaged tumours. Shannon entropy was calculated from mitochondrial variants in HEC116 WT and SMARCA4 KO F0 and F1 tumours to assess clonal heterogeneity. (c) UMAP visualization of Harmony-integrated scRNA-seq data from HEC116 WT and SMARCA4 KO serially passaged tumours, coloured by transcriptional state clusters (C1–C5) and the distribution of WT and KO cells across each cluster. (d) Heatmap of transcription factor enrichment scores from single-cell multiome sequencing in serially passaged tumours. Enrichment scores for selected transcription factors (AUC > 0.7) are shown across transcriptome clusters (C1–C5). Rows indicate clusters, and columns indicate transcription factors. (e) UMAP of hypoxia gene set enrichment in scRNA-seq clusters from HEC116 WT and SMARCA4 KO serially passaged tumours, coloured by enrichment scores. (f) Average expression of BMP4, DLX1, CDH1, and PAX8 for each cluster (C1–C5) identified in Harmony-integrated scRNA-seq data. Dot size reflects the percentage of cells in each cluster expressing the gene, and colour intensity indicates the average expression level. (g) UMAP visualization of Harmony-integrated scATAC-seq data from HEC116 WT and SMARCA4 KO tumours. scATAC-seq cells were mapped onto transcriptional clusters (C1–C5) defined by scRNA-seq, with WT and KO cells highlighted to show genotype distribution across clusters. (h) Clonal composition during serial tumour passaging. Each colour indicates a transcriptome cluster (C1–C5) identified from scRNA-seq analysis of single-cell multiome data. Lines show the proportion of cells in each cluster across F0, F1, and F2 passages for HEC116 WT and SMARCA4 KO tumours. (i) Velocity analysis from serially passaged tumours. Velocity vectors are overlaid on UMAPs of HEC116 WT and SMARCA4 KO tumours, illustrating inferred transcriptional dynamics across F0, F1, and F2 passages. Cells are coloured by inferred temporal order along transcriptional trajectories, ranging from early/progenitor-like state to late/differentiated state, and arrows indicate the predicted direction of transcriptional dynamics between clusters. (j) Dual immunohistochemistry staining of SMARCA4 and E-cadherin in human DDEC and CDX tumours. Top panels: representative human DDEC tumours with SMARCA4 mutations, showing both WD and UD regions. Scale bar: 100 µm. Bottom panels: mouse HEC116 F2 tumours comparing WT and SMARCA4 KO. Scale bar: 200 µm. SMARCA4 is stained in brown (nuclear) and E-cadherin in red (membranous/cytoplasmic). (k) Representative patterns of E-cadherin staining (red) in undifferentiated regions of SMARCA4-mutated tumours. Five different patterns are shown: negative, single-cell, clusters (small or large clusters), patchy, and diffuse. Scale bar: 100 µm. (l,m) E-cadherin expression patterns in SMARCA4-mutated tumours. (l) Representative IHC images showing the different E-cadherin staining patterns observed in undifferentiated regions of SMARCA4-mutated tumour cores. Scale bar: 200 µm. (m) Parts-of-whole plot showing the proportion of tumours exhibiting each E-cadherin pattern based on TMA cores. When multiple patterns were present in a single core, the most prominent pattern was used for classification.

Except for cluster C5 in KO tumours, all clusters express epithelial or lineage-specific genes (e.g. *CDH1*, *PAX8*) (**Fig. 4f; Supplementary Fig. 7**), suggesting that histologically dedifferentiated regions may contain cells that are moving between states, rather than cells that are fully reprogrammed. Given these surprising results, we reasoned that there should exist transitional cells, wherein SMARCA4 is lost within well-differentiated regions, and that seemingly undifferentiated areas should randomly express epithelial lineage markers such as *CDH1*. Accordingly, we examined SMARCA4 (brown) and E-CADHERIN (red) staining in the well and poorly differentiated regions of 4 human DDEC tumours, as well as in 10 cores containing only undifferentiated regions. Our results confirmed that SMARCA4-negative cells could be observed in the well-differentiated regions of all tumours (**Fig. 4j**), and that E-CADHERIN-positive cells could be detected in histologically undifferentiated regions (**Fig. 4j**). We determined that E-CADHERIN positive cells could similarly be detected in morphologically undifferentiated regions of SMARCA4 deficient HEC116-derived xenografts (**Fig. 4k**). Finally, quantification of E-CADHERIN staining across 31 cores of SMARCA4-deficient DDEC, demonstrated that this epithelial protein was distributed heterogeneously within over 77% of undifferentiated regions as single cells, clusters, patches, or diffuse regions (**Fig. 4l,m**). Collectively, these results suggest that SMARCA4 loss does not immediately induce dedifferentiation and that the dedifferentiated histopathology is associated with heterogeneous cell populations variably expressing epithelial markers (e.g. E-CADHERIN) but lacking morphological differentiation.

## DISCUSSION

In this study, we made the surprising discovery that dedifferentiation induced by the loss of SMARCA4 is not governed by the stochastic selection of cells with a stem cell-like fate nor the dedifferentiation of cells into an embryonic-like lineage. Rather, we demonstrate that extreme heterogeneity (the manifestation of an undifferentiated histology) can occur due to the rapid movement between transcriptional states (cell state chaos) and that this is associated with epigenomic instability. Using single-cell sequencing and barcoding, we determined that epigenomic dysfunction and the erratic movement between transcriptional states lead to the histological mimicry of an undifferentiated tissue. Accordingly, cellular states likely switch independently of location within the tumour. This leads to the expression of epithelial-specific proteins (e.g. E-CADHERIN) in otherwise undifferentiated areas. While not a focus of this study, we believe that this phenomenon may not be specific to SMARCA4-deficient DDEC, as we similarly detected E-CADHERIN in the undifferentiation regions of ARID1A/1B-deficient DDECs (**Supplementary Fig. 10)**. Hence, cell state chaos, rather than directional dedifferentiation, may underpin the emergence of histologically undifferentiated cancers.

The KO of SMARCA4 did not immediately cause dedifferentiation in our model, and we could detect SMARCA4-deficient cells within the well-differentiated components of human DDECs (**Fig. 1,2,4j**). Moreover, the loss of SMARCA4 initially induced a senescent-like phenotype, suggesting the existence of a transitional state that could be targeted to prevent disease progression in at-risk endometrial cancer patients. Previous studies have shown that knockdown of SMARCA4 in colorectal cancer cells similarly induces a senescent-like phenotype, characterized by increased β-galactosidase-positive cells and p21 protein^44,45^. In colorectal cancers, this senescent-like phenotype was mediated by the SIRT1/p53/p21 axis^45^. In our HEC116 model, we observed a decrease in p21. However, in the HEC59 model, p21 protein increased upon SMARCA4 loss (**Supplementary Fig. 3d**). Additionally, both cell lines are *TP53* mutants. Hence, while the endometrial cancer cell models used in this study showed reduced sphere and colony formation and increased ß-galactosidase activity, our results suggest that an alternative mechanism for the initiation of a senescent-like phenotype is independent of the p21/p53 axis. Targeting senescent cells has been a promising avenue of therapeutic research in a multitude of cancers. Senolytics target many different pathways involved in senescence, including anti-apoptotic BCL-2 proteins and pro-inflammatory pathways^46^. Interestingly, senomorphics have been investigated to reduce the secretion of identified SASP factors and pro-inflammatory cytokines. For example, metformin has been shown to reduce pro-inflammatory pathways and SASP secretion, specifically IL-6 and IL-8^47^. Further studies should aim to characterize the mechanism underlying the senescent-like phenotype observed in our model to understand how this pathway could be targeted therapeutically in SMARCA4-deficient tumours.

In this study, we show global alterations in DNA methylation in SMARCA4-deficient cells after *in vivo* passaging. This is consistent with human cases of DDEC, which have been associated with hypomethylation^36^. DNA methylation is critical for cell fate specification, and reprogramming toward a pluripotent state is associated with a global reduction in this epigenetic mark. Moreover, studies have shown that alterations in DNA methylation may underpin cancers with dysregulated differentiation. For example, dedifferentiated chondrosarcoma (DDCS), which is a low-grade chondrosarcoma showing high-grade sarcomatous transformation that can display heterologous non-chondroid mesenchymal differentiation (making skeletal muscle, fat, bone), is associated with global hypermethylation^4^. Evidence suggests that *IDH1/2* mutations are responsible for this altered DNA methylation signature in DDCS compared to conventional chondrosarcoma^48^ and that hypermethylation occurs at the promoter regions of *CDH1* and *CDKN2A*^49^. RNA sequencing showed that there is a reduction in *TET1*, *TET2*, *TET3* (that catalyze DNA demethylation), and a reduction and increase in *DNMT3A* and *DNMT3B*, respectively (**Supplementary Fig. 11**) in SMARCA4 KO cells. While this suggests that SMARCA4 deficiency may alter DNA methylation regulation, further studies are warranted to explore this possibility and determine the extent to which alterations in DNA methylation underpin cell state chaos in endometrial cancers. Indeed, the DNA methylation regulatory machinery may be an attractive target for this otherwise fatal disease.

Single-cell multi-omics revealed that SMARCA4 KO tumours contained a small population absent in WT tumours. This cluster was present in early passages and did not increase proportionally with the emergence of dedifferentiated regions during tumour propagation. Moreover, trajectory analyses and lineage tracing suggested that all SMARCA4 KO cells can move in and out of this state, and that it closely associates with other states present in WT cells. Indeed, this population did not express genes specific to any pathway (**Fig. 4**). Hence, we surmise that the BMP4, DLX1-positive population, present only in SMARCA4 KO tumours, occurs randomly (likely due to epigenomic dysregulation) and does not serve as a cancer stem cell for DDEC.

In conclusion, we have demonstrated that SMARCA4 loss can induce DDEC by enabling cell state chaos. This contrasts with lineage plasticity, which is largely directional and occurs in response to cues from the microenvironmental neighbourhood. Hence, epigenetic and transcriptional chaos, rather than directional dedifferentiation, may underpin the emergence of histologically undifferentiated cancers. The dogma is that cancer cells acquire a mutation that causes them to dedifferentiate. Our surprising results challenge this paradigm.

## METHODS

### Cell lines and cell culture

Human endometrial carcinoma HEC116 and HEC59 cell lines were obtained from the Japanese Collection of Research Bioresources (JCRB) Cell Bank and maintained in Minimum Essential Medium (MEM; Gibco) supplemented with 10% fetal bovine serum (FBS; Sigma). Cells were cultured at 37°C in a humidified incubator with 5% CO₂ and passaged using 0.25% (w/v) trypsin (Life Technologies). Authentication of all cell lines was performed at the SickKids Research Institute, and routine mycoplasma testing was conducted in-house using the PCR-based Universal Mycoplasma Detection Kit (ATCC).

### CRISPR/Cas9-derived SMARCA4 knockout and clone validation

HEC116 and HEC59 cells were edited using either a plasmid- or ribonucleoprotein (RNP)-based CRISPR/Cas9 system. For the plasmid-based approach, a custom gRNA (sequence: CGCCCGTGATGCCACCGC) was cloned into an all-in-one CRISPR/Cas9 LacZ plasmid and transfected into HEC116 cells using GeneIn™ (GlobalStem). mCherry⁺ cells were enriched by flow cytometry, and single clones were generated by sorting into 96-well plates. For the RNP approach, crRNA (sequence: GCGGTGGCATCACGGGCG) and tracrRNA were annealed at 95°C, followed by gradual cooling, complexed with Alt-R *S. pyogenes* Cas9 (IDT) in Opti-MEM (Gibco) for 5 min, then combined with Lipofectamine RNAiMAX (Invitrogen) and applied to cells (4×10⁴ cells/well; 10 nM RNP final). tracrRNA-ATTO550⁺ cells were enriched by flow cytometry and plated as above. Genomic DNA was extracted from HEC116 and HEC59 colonies using PureLink Genomic DNA Isolation Kit (Invitrogen) according to the manufacturer’s instructions and quantified by spectrophotometry. Single-cell-derived clones were validated by Sanger sequencing of the respective target region. PCR amplification was performed with AmpliTaq Gold (Applied Biosystems), and PCR products were cloned using the TOPO TA Cloning Kit (Invitrogen). Bacterial colonies were sequenced using M13R primer targeting the pCR-4-TOPO vector backbone to confirm editing of all alleles.

### Animal Studies

All animal experiments were performed in compliance with the Canadian Council on Animal Care guidelines and approved by the University of Alberta Animal Policy and Welfare Committee. Cell line-derived xenograft (CDX) tumour fragments (approximately 2 mm in diameter) from HEC116 plasmid system-derived (PD) or HEC59 RNP system-derived WT or SMARCA4 KO tumours expanded via serial passaging and collected at the third generation, were implanted subcutaneously into 6–8-week-old female NOD.Cg-Prkdc^scidIl2rg^null (NSG) mice. Fourteen days after implantation, animals were randomized into treatment cohorts receiving either vehicle control (phosphate-buffered saline; PBS) or carboplatin (Cayman Chemicals) at a dose of 60 mg/kg administered intravenously once a week. Treatment was continued until tumours reached a maximum diameter of approximately 10 mm. Tumour length and width were measured twice weekly using digital callipers, and tumour area was calculated for growth analysis. Tumour growth rates were calculated as the slope of a linear regression of the natural log of tumour volume over time, and statistical analysis was performed using multiple t-test with Holm–Šídák correction in GraphPad Prism. At the experimental endpoint, defined as a tumour area of approximately 60 mm², mice were euthanized, and tumours were excised and weighed.

### Xenograft formation, tumour dissociation, and *in vivo* serial passaging

For xenograft establishment, 3×10⁶ cells were resuspended in 100 μL of MEM and Matrigel (Corning) at a 1:1 ratio and injected subcutaneously into both the left and right flanks of 7–8-week-old female NSG mice. When tumours reached approximately 10 mm in diameter, each was bisected: one half was fixed in formalin and paraffin-embedded for histological and immunohistochemical analyses (University of Alberta Laboratory Medicine and Pathology Core Facility), and the other half was either flash frozen and stored at −80°C for subsequent RNA extraction, or subjected to mechanical and enzymatic dissociation using the MACS Human tumour Dissociation Kit (Miltenyi Biotec). The protocol was followed according to the manufacturer’s instructions, with the sole modification of omitting the final strainer wash. Dissociation was performed using the gentleMACS Dissociator (Miltenyi Biotec) with the “37C_h_TDK_3” program optimized for fibrous tumour tissue. Single-cell suspensions obtained from dissociated tumours were reinjected into recipient mice following the same procedure described above for serial passaging.

### Immunohistochemistry for cell lines and tissues

Formalin-fixed paraffin-embedded (FFPE) tumour tissues and agarose-embedded cell pellets were processed for immunohistochemical staining. Cells in culture were pelleted, fixed overnight in 4% paraformaldehyde (PFA; Chem Cruz), and embedded in 1% agarose before paraffin embedding. Sections were deparaffinized in xylene, rehydrated through a graded ethanol series, and subjected to antigen retrieval using pH 9 buffer (Dako) for general staining or Cell Conditioning Solution (CC1; Tris-based EDTA buffer, pH 8, Ventana) for PAX8 detection. Endogenous peroxidase activity was quenched with peroxidase block for 5 minutes, followed by serum-free protein blocking for 10 minutes. Primary antibodies were applied for either 30 minutes at 25°C or overnight at 4°C for manual staining, or according to the protocols for the Ventana Discovery XT, Benchmark XT, and Benchmark Ultra automated systems. Antibodies and working dilutions were as follows: anti-BRG1 [EPNCIR111A] (Abcam, 1:200 for manual staining, and 1:25 for automated Ventana staining), anti-PAX8 BC12 [ACI 438] (Biocare Medical; 1:100), and E-cadherin G10 [sc-8426] (Santa Cruz Biotechnology; 1:100). For automated PAX8 staining, primary antibody incubation was performed for 1 hour at 37°C with Ventana antibody diluents, followed by Ventana Universal Secondary Antibody for 25 minutes at 37°C. For manual staining, Envision HRP anti-rabbit or anti-mouse secondary antibodies (Dako) were incubated for 1 hour at room temperature. Detection was performed with DAB substrate (Dako), followed by counterstaining with Mayer’s hematoxylin (Vector Laboratories), dehydration through a reverse ethanol series, and coverslipping with Vectamount mounting medium (Vector Laboratories). Whole-slide imaging was performed using an Aperio Digital Pathology slide scanner, and image analysis was conducted using ImageScope software (Leica Biosystems). Staining images from human specimens were provided by Dr. Cheng Han-Lee. Mice slides were stained either on the Bond III automated system (Leica) at Kingston Health Sciences Centre or on Ventana Discovery/Benchmark platforms (University of Alberta). For Bond III, slides were deparaffinized with Bond Dewax Solution (Leica) and subjected to heat-induced epitope retrieval with BOND ER Solution 2 (EDTA-based, pH 9). Endogenous peroxidase activity was quenched with Refine Detection Kit Peroxide Block for 5 minutes, followed by serum-free protein blocking for 20 minutes. Sections were incubated with anti-BRG1 [EPNCIR111A] (Abcam; 1:500) for 15 minutes at room temperature and anti-E-cadherin [NCH-38] (Dako; 1:200) for 15 minutes at room temperature. Secondary detection was performed with Refine Detection Kit Polymer for 15 minutes at room temperature, and antigens were revealed with DAB for BRG1 (nuclear staining) and Fast Red for E-cadherin (cell membrane/cytoplasm staining). Slides were counterstained with Refine Detection Kit Hematoxylin (Leica), dehydrated through a graded ethanol series, and coverslipped using the Leica CV5030 with Permount mounting medium (Fisher Scientific). For Ventana-stained mouse slides, SMARCA4 and E-cadherin staining was performed with pH 9 HIER and DAB detection, followed by hematoxylin counterstaining and scanning using the Aperio Digital Pathology system. Cell pellets were manually stained for SMARCA4 under the same conditions as above. Dual SMARCA4/E-cadherin staining was performed on a tissue microarray (TMA) provided by Dr. Cheng Han-Lee and stained on the Bond III automated system at Kingston Health Sciences Centre as described, with DAB for BRG1 and Fast Red for E-cadherin, followed by hematoxylin counterstaining, dehydration, and coverslipping.

### RNA extraction, library preparation, sequencing and analysis

RNA was extracted from serum-starved cells using the RNeasy Kit (Qiagen) and quantified with the Qubit RNA HS Assay Kit (Invitrogen). Samples were sent to the Genome Quebec Innovation Center for quality assessment using the 2100 Bioanalyzer (Agilent), followed by library preparation and sequencing. Stranded mRNA libraries were prepared with the NEBNext Ultra II Directional RNA Library Prep Kit for Illumina (NEB) and sequenced on the Illumina NovaSeq 6000 platform. Raw reads were assessed for quality using FastQC (default parameters). Adapters and low-quality reads were removed with trimmomatic (paired-end mode). Filtered reads were aligned to Homo sapiens reference genome GRCh38 (Ensembl release #106) using HISAT2. Mapped reads were quality-checked with samstat, sorted and indexed with samtools, and further processed with Picard. Gene-level counts were generated using featureCounts. Differential expression analysis was performed in R using the DESeq2 package, and significantly differentially expressed genes (FDR < 0.01 and log₂FC > |1|) were subjected to functional enrichment analysis using the DAVID database (updated April 2023) and Gene Set Enrichment Analysis (GSEA).

### Quantitative Real-Time PCR (qPCR)

Total RNA was extracted from cultured cells using RNeasy Kit (Qiagen) with on-column DNase treatment and eluted in 40 µL RNase-free water. RNA concentration and purity were measured using an Epoch Microplate Spectrophotometer (BioTek). Complementary DNA (cDNA) was synthesized from 2 µg of total RNA using the High-Capacity cDNA Reverse Transcription Kit (Applied Biosystems) with a 1:1 mixture of random hexamers and oligo(dT) primers (Invitrogen). No-template and no-reverse transcriptase controls were included. qPCR reactions were performed using 1 µL of cDNA, TaqMan Gene Expression Master Mix (Applied Biosystems), and FAM-labeled TaqMan Gene Expression Assays for *CDH1* (Hs_01023894_m1), *CLDN3* (Hs00265816_s1), *OCT4* (Hs_04260367_gH), and *ESR1* (Hs_00174860_m1). Amplification was performed on a CFX96 Touch Real-Time PCR Detection System (Bio-Rad) with the following thermocycling conditions: 95°C for 10 min, followed by 40 cycles of 95°C for 15 s and 60°C for 1 min. Relative mRNA expression levels were calculated using the ΔCT method, comparing SMARCA4 KO to control samples.

### NanoString gene expression profiling

Gene expression profiling was performed on five matched FFPE DDEC tissue samples obtained from Calgary Laboratory Services and the Royal Alexandra Hospital, as well as on serially passaged CDX tumours from NSG mice. The PanCancer Progression Panel (NanoString), containing 770 genes including 30 reference genes associated with epithelial–mesenchymal transition, extracellular matrix remodelling, metastasis, and angiogenesis, was used for all samples. RNA was extracted from the undifferentiated and differentiated components of human tumour cores (3–6 cores each) and murine CDX tumours, following appropriate deparaffinization using the FFPE RNA Extraction Kit (Roche). RNA quality and concentration were assessed using 2100 Bioanalyzer (Agilent), and 100 ng of total RNA was hybridized with fluorescently barcoded 3′ biotinylated capture probes and 5′ reporter probes for each target at 65°C overnight. Samples were processed on the nCounter Digital Analyzer (NanoString) for direct transcript counting. Raw data were analyzed using nSolver Analysis Software v4.0 (NanoString). Background signal was determined from the mean count of eight negative control probes per reaction. Counts were normalized to the geometric mean of six internal positive controls. Fold-change estimation between undifferentiated (sample) and differentiated (reference) components was performed using two-tailed Student’s t-tests when cases were pooled as replicates. For individual case analyses, differential expression was determined based on NanoString confidence limits at each expression count level.

### Tumoursphere formation assay

Single-cell suspensions were obtained by filtering the suspension through a 40 μm cell strainer before counting viable cells using trypan blue. Cells were diluted to the desired concentration in sphere media composed of MEM supplemented with 1x B27 (Gibco), 20 ng/mL EGF (Sigma), and 10 ng/mL FGF (Sigma). 100 μL of the cell suspension was plated per well in 96-well Ultra Low Attachment (ULA) plates. Cultures were maintained at 37°C and 5% CO₂ for 14 days. Spheres were manually counted under a light microscope. Representative images were captured using EVOS FL Cell Imaging System (Thermo Scientific). Statistical analysis was conducted using multiple t-tests with Holm-Šidák correction in GraphPad Prism.

### Soft agar colony formation assay

Six-well plates were pre-coated with a 1:1 mixture of 1% agarose and 2x MEM and 1,000 cells were embedded in a top layer containing 0.7% agarose mixed 1:1 with 2X MEM. After 14 days, colonies were washed with PBS and stained with 0.5% crystal violet solution. Colonies were manually counted under a light microscope. Representative images were captured using EVOS FL Cell Imaging System (Thermo Scientific). Statistical analysis was performed using an unpaired t-test in GraphPad Prism.

### β-Galactosidase senescence assay

For senescence-associated β-galactosidase (SA-β-Gal) detection, cells were either seeded onto glass coverslips and placed in 6-well plates or directly seeded into 6-well plates. The next day complete growth medium was removed, and cells were rinsed with 10 mL of pre-warmed PBS. To minimize serum contamination, serum-free medium was added after the PBS wash, and cells were incubated for 1 hour at 37°C. Subsequently, 20 mL of basal medium was added for 24 hours. Cells maintained in complete growth medium were used as negative controls. Cells were then fixed in 1 mL of Fixative Solution (Senescence β-Galactosidase Staining Kit; Cell Signaling Technology) for 10–15 minutes at room temperature, rinsed twice with PBS, and incubated with 1 mL of β-Galactosidase Staining Solution per well, according to the manufacturer’s protocol. Plates were sealed and incubated overnight at 37°C in a dry incubator. Following staining, the solution was removed, and coverslips were mounted onto glass slides, and plates were stored in 70% glycerol. Brightfield images were acquired using a Zeiss Axioskop microscope (University of Alberta, Cross Cancer Institute, Imaging Facility). Five random fields of view per well were imaged, and the total number of cells and β-Gal–positive cells were quantified using ImageJ. The percentage of senescent cells was determined as the ratio of blue-stained cells to the total cell count. Statistical analysis was performed using an unpaired t-test in GraphPad Prism.

### H3K9me3 immunostaining

Cells adherent to coverslips were fixed with 4% PFA for 10 minutes and permeabilized with 0.5% Triton X-100 for 5 minutes. Primary αH3K9me3 antibody (Active Motif; 1:1000) was incubated for 30 minutes at RT, followed by one wash with 0.1% Triton X-100 for 1 minute and three rinses with 1X PBS. Secondary antibody (Goat anti-rabbit Alexa Fluor 488; 1:500) was incubated for 30 minutes at RT and washed as above. Coverslips were mounted on microscope slides using PVA mounting media (Supelco) containing 1 μg/mL DAPI (Invitrogen). Images were acquired on a Zeiss LSM 710 Meta confocal microscope with the pinhole set to 1 airy unit for all channels and exposure gains kept constant across samples. Confocal images were analyzed using Imaris v9.2.1. Statistical analysis was performed using an unpaired t-test in GraphPad Prism.

### Protein extraction and Western Blot

Protein lysates were prepared from adherent cells directly on culture plates using radioimmunoprecipitation assay (RIPA) Lysis and Extraction Buffer (Thermo Scientific) supplemented with 1% Halt™ Protease and Phosphatase Inhibitor Cocktail (Thermo Scientific). Lysates were incubated on ice for 15 minutes and centrifuged at 13,000 rpm for 20 minutes at 4°C to remove insoluble debris. Soluble protein concentrations were determined using the DC™ Protein Assay (Bio-Rad) and measured with a FLUOstar Omega plate reader (BMG LABTECH). Equal amounts of protein were combined with 4X Laemmli sample buffer (Bio-Rad) containing 5% (v/v) 2-mercaptoethanol (Sigma), boiled for 5 minutes, and resolved on 7.5%-12% SDS-PAGE gels. Proteins were transferred to nitrocellulose membranes (Bio-Rad) at 100 V for 90 minutes. Membranes were blocked in 5% (w/v) non-fat dry milk in TBST for 1 hour at room temperature and subsequently incubated overnight at 4°C with the following primary antibodies: p21 [EPR3993] (Abcam; 1:500), H3K9me3 [D4W1U] (Cell Signaling Technology; 1:1000), or β-Tubulin (LI-COR; 1:7000). After washing in TBST, membranes were incubated for 1 hour at room temperature with horseradish peroxidase (HRP)-conjugated secondary antibodies (Bio-Rad; 1:3000). Detection was performed using Clarity™ Western ECL Substrate (Bio-Rad), and images were captured on Fujifilm X-ray film. Band intensities were quantified by densitometry using ImageJ.

### Mass spectrometry-based proteome

Sample preparation: HEC116 and HEC59 cells were plated into T150 flasks or 100 mm plates and cultured to 70–90% confluency. For serum depletion, growth medium was replaced with serum-free medium after rinsing cells with 10 mL of pre-warmed PBS. Conditioned media (CM) was collected, centrifuged at 450 g for 5 min at 4°C, and stored at −80°C until processing. For whole-cell proteome analysis, cells were lysed in RIPA buffer supplemented with 1% Halt™ Protease and Phosphatase Inhibitor Cocktail. Lysates were incubated for 15 minutes, sonicated, and centrifuged at 14,000 g for 15 minutes at 4°C to remove debris. Supernatants were stored at −80°C until further processing.

HEC59 CM processing: Thawed CM samples were concentrated using 3 kDa MWCO ultra-centrifugal units (Millipore) at 4000 g for 45 min at 8°C. Salts and residual medium were removed by adding 10 mL of distilled water to the filter unit, followed by centrifugation for 15 minutes at 4000 g. The final concentrated volume (500 μL) was stored at −80°C, lyophilized overnight, and resuspended in urea-based lysis buffer (8 M urea, 2% SDS, 10 mM DTT, 5 mM ammonium bicarbonate). Protein concentration was determined using the Pierce™ 660 nm protein assay with ionic detergent compatibility reagent. Samples were incubated for 30 minutes at RT with 10mM DTT and alkylated with 100 mM iodoacetamide (IAA) for 30 minutes in the dark. Proteins were precipitated using methanol–chloroform extraction, resuspended in 50 mM ammonium bicarbonate, and sonicated. Digestion was performed with Trypsin/LysC (Promega) at a 1:50 enzyme:protein ratio overnight at 37°C, followed by a second digestion at 1:100 for 4 h. Digestions were carried out at 400 rpm overnight and 1400 rpm for 4 h. Reactions were acidified with 10% formic acid and centrifuged at 14,000 g for 1 min to remove insoluble material.

HEC116 CM and cell lysate processing: Lyophilized CM or cell pellets were lysed in 6 M urea, 2% SDS, 1% NP-40, 0.1% DDM, 50 mM ammonium bicarbonate, followed by reduction with 5 mM TCEP and alkylation with 25 mM IAA for 30 minutes in the dark. Proteins were precipitated using the SP3 protocol^50^ with slight adaptations: precipitation in 70% acetonitrile, followed by three washes with 80% ethanol and a final wash with 95% acetonitrile. Pellets were resuspended in 0.01% DDM and 50 mM ammonium bicarbonate. Proteolysis was carried out with LysC (1:100, 3h, 37°C, 900 rpm), followed by trypsin (1:50, 18h, 37°C, 900 rpm). Peptides were captured with water and acidified with 1% TFA, desalted on in-house C18 StageTips, washed with 0.1% formic acid, and eluted with 60% acetonitrile/0.1% FA. Eluted peptides were dried by vacuum centrifugation at 45°C for 1–2 h and resuspended in 0.01% DDM/0.1% FA.

LC-MS/MS analysis: HEC59 peptides (1 μg) were analyzed on a Q Exactive Plus (ThermoFisher Scientific) coupled to a Waters ACQUITY M-Class UPLC. Samples were loaded onto a Symmetry C18 trap column and trapped for 6 min at 5 μL/min in 99% Solvent A (water/0.1% FA) and 1% Solvent B (acetonitrile/0.1% FA), followed by separation on a Peptide BEH C18 column at 300 nL/min, 35°C, using a non-linear gradient: 1–7% B (1 min), 7–23% B (179 min), 23–35% B (60 min). MS settings followed as previously described^51,52^. HEC116 peptides (200 ng) were analyzed on a Vanquish Neo coupled to a Thermo Eclipse mass spectrometer. Samples were loaded onto a trap column and separated on a 25 cm IonOpticks Aurora Series column at 50°C using a non-linear gradient: 2–5% B (2 min), 5–40% B (50 min), 40–60% B (5 min), and 60–95% B (3 min).

Gas Phase Fractionation Data-Independent Acquisition (GPF-DIA): For spectral library generation, a pooled digests of representative CM and whole-cell lysate samples was serially injected to produce 100 m/z fractions across 400–1000 m/z using a staggered 4m/z window scheme, resulting in 2 m/z bins after demultiplexing^53^. CM or cell lysate peptides (1 μg) were analyzed using staggered 24 m/z windows, producing 12 bins after demultiplexing. Raw files were converted to mzML using ProteoWizard (PeakPicking=1, Demultiplex=10 ppm, ZeroSamples=−1) and searched together with the spectral library in DIA-NN, allowing two missed cleavages, one variable oxidation, and fixed carbamidomethylation on cysteine.

Data analysis: Label-free quantification (LFQ) data were processed in R using the DEP package. Background correction and variance stabilizing transformation (VST) were applied, followed by MinProb imputation of missing values. Functional enrichment of significantly altered proteins (p < 0.05, |log₂FC| > 0.5) was performed using DAVID.

### DNA extraction and array-based methylation analysis

Genomic DNA was extracted with PureLink Genomic DNA Isolation Kit (Invitrogen) following manufacturer’s instructions. DNA concentration was determined by spectrophotometry using Epoch Microplate Spectrophotometer (BioTek). Samples were analyzed using the Illumina Infinium EPIC (850 k) BeadChip (Illumina). Data processing and analysis were performed in R using the minfi package, with preprocessing steps including background correction and dye-bias normalization. Probes located on sex chromosomes, those containing multiple single-nucleotide polymorphisms (SNPs), and probes lacking unique genomic mapping were excluded from further analysis. Pairwise sample relationships were visualized using t-distributed stochastic neighbour embedding (t-SNE) applied to the 10,000 most variable probes using perplexity of 3 and 3,000 iterations. Group comparisons were conducted to identify differentially methylated positions (DMPs) and differentially methylated regions (DMRs) based on M-values using the empirical Bayes (eBayes) method. Probe annotations were assigned according to the Illumina Human MethylationEPIC array manifest (Epic.ilm10b4.hg19)^54^.

### Whole Exome Sequencing and analysis

Genomic DNA was extracted from unpassaged cells and second generation (F2) *in vivo* serially passaged tumours using the PureLink Genomic DNA Isolation Kit (Invitrogen) according to the manufacturer’s instructions. Purified gDNA was submitted to Genome Quebec Innovation Center for library construction, quality control, exome enrichment, and sequencing. Whole-exome libraries were prepared using the SureSelect Human All Exon V7 kit (Agilent) for target enrichment. Paired-end sequencing was performed on an Illumina NovaSeq 6000 platform, generating ∼25 million reads per sample. All samples were processed and sequenced in a single batch to avoid technical batch effects. Somatic variants were called using Mutect2 (v. 4.6.2) and subsequently filtered with FilterMutectCalls to retain only those labelled “PASS.” Germline variants were called using HaplotypeCaller (v. 4.6.2). Both somatic and germline variants were normalized and filtered using BCFtools before annotation with SnpEff (v. 5.2). To enable comparisons across tumour microenvironments and to account for CRISPR-induced mutations that are detected as germline rather than somatic by variant callers, both germline and somatic variants were analyzed. For downstream analyses, only nonsynonymous variants located in cancer-related genes were included. Cancer-related genes were defined according to the COSMIC Cancer Gene Census (v.102, released May 21, 2025). Tumour mutational burden (TMB) was calculated as the number of somatic, coding base substitutions and indels per megabase of the genome covered by at least 10 reads. Heatmaps were generated using the allele frequency of nonsynonymous variants in cancer-related genes across samples.

### Lentiviral barcoding

Barcode oligonucleotides were designed using forward (5’-TCGAGAAGTAA NNATCNNGATSSAAANNGGTNNAACNNTGTAAAACGACGGCCAGTGAGC-3’) and reverse (5’- CCGGGCTCACTGGCCGTCGTTTTACANNGTTNNACCNNTTTSSATCNNGATNNTTACT TC-3’) sequences, annealed, and ligated into an MPG vector downstream of a GFP reporter. Bacteria were transformed and plated at clonal density onto ampicillin agar plates. Colonies were pooled and purified following the Plasmid Maxi Kit (QIAGEN) manufacturer’s instructions. Single colonies were validated by Sanger sequencing to ensure barcoding diversity prior to lentiviral production^22^. Lentiviral particles were produced in HEK293T cells cultured in Dulbecco’s Modified Eagle Medium (DMEM) supplemented with 10% FBS, 4 mM L-glutamine, 1 mM sodium pyruvate, and 0.1 mM non-essential amino acids, using third-generation packaging plasmids (pRRE, pREV, pVSVG) and Lipofectamine-mediated transfection. Viral supernatants were collected, centrifuged, and filtered through a 0.45 µm syringe filter. Cells were transduced with viral supernatant supplemented with 8 µg/mL polybrene. GFP-positive cells were identified and sorted by flow cytometry within the Flow Cytometry Core (RRID:SCR_019195) at the University of Alberta (yields: 18.6% GFP+ in WT and 22.1% GFP+ in KO). Sorted cells were replated and expanded for downstream lineage tracing experiments. Genomic DNA was extracted using a PrepGEM DNA extraction kit (ZyGEM) and barcode amplicons were generated via a 35-cycle PCR reaction using Illumina MiSeq-compatible primers. Libraries were column-purified, quality-checked, and sequenced by the Advanced Cell Exploration Core (RRID:SCR_019182) on a 150-cycle MiSeq kit with 40% phiX spiked in to improve cluster recognition^22^.

### Single-cell multiome RNA and ATAC sequencing

Tumour cells from serially passaged CDX were dissociated and prepared as described in “Xenograft formation, tumour dissociation, and *in vivo* serial passaging”. Mouse and dead cells were removed using the MACS Mouse Cell Depletion Kit and Dead Cell Removal Kit (Miltenyi Biotec). Nuclei isolation and library preparation followed the 10x Genomics Chromium Next GEM Single Cell Multiome ATAC + Gene Expression protocol. Briefly, 500,000 cells were lysed on ice for 3 minutes with Lysis Buffer (10 mM Tris-HCl pH 7.4, 10 mM NaCl, 3 mM MgCl2, 0.1% Tween-20, 0.1% Nonidet P40 Substitute, 0.01% digitonin, 1% BSA) and resuspended in Diluted Nuclei Buffer (10X Genomics) at 5000 nuclei/μL. scATAC-seq libraries were prepared following the Chromium Single Cell ATAC Reagent Kits User Guide (10X Genomics). Nuclei were incubated with the transposase mix to fragment DNA and add the adapter sequences, followed by generation of the gel bead-in emulsions, nuclei barcoding, and addition of Illumina P5 (Read 1N). Illumina P7 and sample index were added during library construction via PCR. All the steps followed the manufacturer’s protocol, using the Chromium Controller (PN-1000202), Chromium Next GEM Single Cell ATAC Library & Gel Bead Kit v1.1 (PN-1000176), the Chromium Next GEM Chip H Single Cell Kit (PN-1000162) and the Single Index Plate N Set A (PN-3000427). Libraries were sequenced on Illumina NovaSeq at ∼25,000 read pairs per nucleus for gene expression and ATAC (∼50,000 total reads) (Novogene, Sacramento, CA, US). Data were processed using the CellRanger ARC pipeline and analyzed using Seurat v5.2.0 and ArchR v1.0.3. DoubletFinder v2.0.6 was used to remove doublets from the dataset^55^. Low-quality scRNA-seq transcripts were filtered (counts >25,000 and <=1,000; features >7,500 and <=800; mitochondrial >25%; ribosomal >10%, and hemoglobin >0.05), and samples were integrated using Harmony. 10 PCA dimensions were used to find nearest neighbours, and clusters were generated with a resolution of 0.2. scATAC-seq data were processed with ArchR’s multiome analysis pipeline (hg38), and clusters were mapped to the Seurat object^56^. All single-cell work was conducted at the Advanced Cell Exploration Core, University of Alberta (RRID: SCR_019182).

### Transcription Factor and Pathway Analysis

To infer regulatory programs, we applied the decoupleR framework to the log-normalized “RNA” assay of the integrated Seurat object. Transcription factor activity was estimated with the human CollecTRI regulatory network, which links each factor to its experimentally supported target genes and specifies the direction of regulation^57^. For each biological sample, we extracted the gene expression matrix for the corresponding cells and scored regulons using the weighted mean method implemented in run_wmean. The procedure permuted target labels one hundred times to generate a background distribution and reported the normalized weighted mean score. Regulators with fewer than 5 target genes were ignored. Pathway activity was computed in an analogous manner using the human PROGENy database^58^, restricted to the 100 most responsive genes for each pathway. Weighted mean scoring with one hundred permutations and a minimum target set size of five produced normalized activity values for every pathway in every cell. For each assay, we used FindAllMarkers with the receiver-operating characteristic (ROC) classifier, which ranks features by how well their values separate a given cluster from all other cells.

### Velocity analysis

Latent time and velocity analysis were performed using the scvelo4 package pipeline^43^.

### Mitochondrial lineage tracing

As a first step, we generated per position coverage and variant read count information across the mitochondrial genome. To do so, we used a pipeline modified^59^. This involved first splitting the reads aligned to the mitochondria (“chrM”) for each retained cell barcode into separate files (using ‘01_split_bam_by_barcode_v2’). We next generated per-position total coverage and counts of each base, including only reads with a minimum Phred base quality threshold of 23.8 and a minimum alignment quality of 30 (using ‘02_pileup_counts_bulk.py’). Individual per position base and coverage counts were then combined into a single R ‘MultiAssayExperiment’ object for downstream analysis (using ‘03_makeRDSv4.R’). This was done separately for the BAM files from the ATAC and RNA data generated by Cell Ranger. An example workflow is as follows:

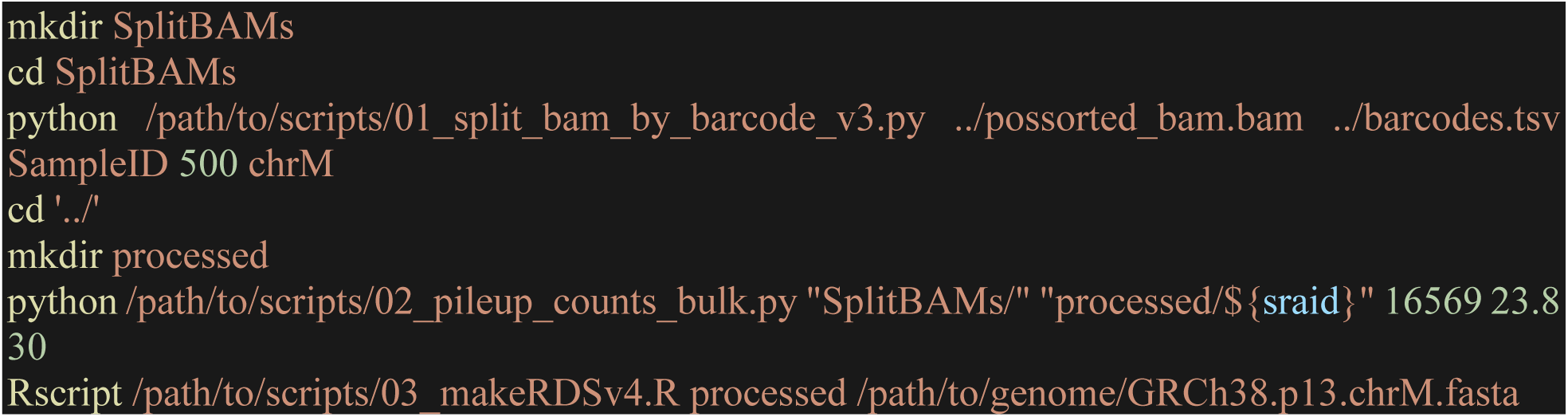

For each of the six passage–genotype samples, we loaded the per-cell mitochondrial mutation count and coverage matrices derived separately from the matched scRNA-seq and scATAC-seq libraries. Cells whose barcodes were present in both modalities were identified, and for every shared barcode, the allele-specific counts and coverages were summed, creating a single modality agnostic estimate per cell and genomic position; a sanity check was performed to confirm that summed counts never exceeded total coverage.

To construct a common coordinate system, we next restricted the analysis to alleles observed in all six samples, concatenated the individual matrices column-wise, and verified once more that counts did not exceed coverage. Sites with fewer than five mutant reads in at least five cells and cells that were not retained in the multiome analysis pipeline were discarded, yielding a pooled object containing a mutant count matrix and the corresponding coverage matrix.

Maximally informative positions and cells were then selected with a two-stage, data-driven elbow procedure. Each matrix entry was first assigned a confidence weight by passing its read depth through a logistic curve with a midpoint of 30 reads and a slope parameter of 0.1; this transformation yields values close to one for well-covered positions and values close to zero for very shallow coverage. The mutant-allele frequency, defined as the number of mutant reads divided by the total reads at that position, was then multiplied by its corresponding weight to create an information-score matrix. We gauged the informativeness of each genomic position by calculating the variance of its information scores across all cells, ranked the positions from most to least variable, and retained those above the elbow point of the resulting curve (the rank at which the curve bends most sharply away from a straight line). Using only those positions, we added up the information scores within each cell, ranked the cells, applied the same elbow rule, and retained the cells above that second elbow.

Distances between the retained cells were computed by recalculating allele frequencies on the filtered matrix and then, for every pair of cells, taking the squared difference in allele frequency at each jointly covered informative site, multiplying it by the minimum of the two site-specific weights, summing across sites, and normalizing by the total of those minimum weights. The square root of this normalized sum yielded a coverage-weighted Euclidean distance. Pairs sharing no jointly covered informative sites were assigned NA. This weighted distance matrix was used for downstream calculations of clonal distribution.

### Clonal Clustering and Entropy

To objectively identify clonal groups, we converted the weighted distance matrix into a K-Nearest Neighbour graph (with k = 20), treating smaller genetic distances as stronger connections. Community detection was then performed using the Leiden algorithm with the R package ‘leiden’, while sweeping the resolution parameter from 0.1 to 1.5 in increments of 0.1. For each resolution, we generated a two-dimensional Uniform Manifold Approximation and Projection (UMAP) embedding (using the R package ‘umap’) that relied on the original distances and colored each point (cell) by its Leiden label. These diagnostic plots were saved to enable visual comparison of clustering granularity. After inspecting these plots, we chose the partition at a resolution of 1.5 because it separated the main genetic lineages without excessive fragmentation. The resulting clusters were assigned distinct, automatically generated colours and used as the working clone labels.

Two complementary low-dimensional representations were created for illustration: the UMAP generated during clustering and a classical metric multidimensional scaling (MDS) projection of the same distance matrix. Using the final clone calls, we quantified clonal diversity within each biological sample. For every sample, we quantified the number of cells assigned to each clone and converted that frequency distribution into a Shannon entropy score using the R package ‘entropy’, providing a single measure of within-sample clonal diversity. To gauge the uncertainty associated with unequal cell numbers, we also performed a bootstrap procedure in which 100 cells were randomly drawn from each sample 1000 times, with replacement, and their clone entropies recalculated.

### Clone molecular composition over time

To compare mitochondrial clones with cell states obtained from the integrated multiome analysis, we first copied the final clone labels into the Seurat object, then concatenated each label with the cell’s sample identifier to generate a unique “clone | genotype” tag. A two-way contingency table was then created with clones as rows and transcriptome-defined clusters as columns. For each clone, we tested whether its distribution across clusters differed from the overall cluster frequencies. We used a Fisher’s exact test with Monte Carlo simulation to calculate a P-value, and the resulting list was corrected for multiple testing using the Benjamini–Hochberg method.

To gauge similarity between clones from a transcriptional perspective, we converted the contingency table into clone-specific cluster-proportion profiles, centred each profile by subtracting the overall cluster frequencies, and projected the clones into two dimensions using metric multidimensional scaling (MDS). The same distance matrix was subjected to average-linkage hierarchical clustering, and the dendrogram was cut at a height of 0.25 to define broad clone groups. Bar-plot panels were generated for every clone, displaying either the raw deviations or the log-fold deviations from the overall cluster composition, so that over- and under-representation of individual RNA states could be inspected at a glance. The MDS and dendrogram were exported with clone labels colored by their tree-based group. A comma-separated file summarized each clone’s group assignment, Fisher P-value, adjusted P-value, and cluster proportions.

For the temporal analysis, we tallied the distribution of cells across RNA clusters within each clone, separately for passages F0, F1, and F2, normalized those counts to proportions, and stored the resulting three-by-five matrices. Pairwise Euclidean distances between clones were computed on these time-resolved profiles, followed by MDS and hierarchical clustering with a cut height of 0.5 to group clones that shared similar clonal trajectories. A stacked-area plotting function rendered, for every clone, the passage-by-passage change in cluster composition, using the same cluster colour scheme; axes and labels were added automatically. The time-aware MDS plot and the grid of stacked-area charts provide an overview of how individual mitochondrial lineages diversify or remain stable in transcriptional state space over time.

### Barcode sequence analysis

FASTQ files were processed using the ‘ShortRead’ package in R to identify and extract barcode sequences from high-throughput sequencing data. Reads containing the barcode sequence (TAANNATCNNGATSSAAANNGGTNNAACNNTGT) without mismatches in the constant region and where variable bases had a Phred score of 30 or higher were retained. Unique barcode sequences were counted, and barcodes with sequence similarity within a Levenshtein distance of 2 were grouped to account for potential sequencing errors. Processed barcode counts and grouped barcode data were output as text files for downstream analysis.

Barcode frequency distributions were analyzed for each sample by calculating the fractional read value (FRV) of each barcode, defined as the proportion of total reads attributed to a given barcode. A minimum of 100 reads was required to be considered in subsequent analyses. Shannon entropy was calculated using the R package ‘entropy’ for each sample to quantify clonal diversity. To assess the robustness of these entropy estimates, a resampling-based approach was applied, with 1000 iterations of random sampling, each selecting 100 barcodes with replacement according to their FRV-derived probability. The entropy of each resampled dataset was calculated, and the median, 5th, and 95th percentiles of the resampled entropies were used to generate confidence intervals.

Pairwise comparisons of barcode presence were performed across samples to determine the number of shared barcodes, and a matrix of shared barcode counts was generated. Histograms were constructed to visualize the distribution of barcode frequencies, with log10-transformed FRV values plotted to facilitate interpretation. Shannon entropy was calculated to assess clonal diversity, with additional resampling-based entropy estimates obtained by bootstrapping barcode distributions 1,000 times. A barplot of the number of clones above the detection threshold was generated for each sample to compare clonal abundances. Venn diagrams were created to illustrate the overlap among barcodes across samples, and a Pearson correlation coefficient was computed to quantify the similarity in barcode abundances. A Kolmogorov-Smirnov test was conducted to determine whether barcode frequency distributions differed significantly between samples. All analyses were performed in R, and visualizations were exported as PDF files for downstream interpretation.

## CODE AVAILABILITY

Scripts for performing mitochondrial analysis processing are available through GitHub at: https://github.com/djhfknapp/Coverage-Weighted-Mito-Dist/tree/main.

## DATA AVAILABILITY

Raw and processed bulk RNA-sequencing (GSE305733), DNA methylation (GSE306526) and single-cell multiome RNA-seq and ATAC-seq (GSE305749) are available on the National Center for Biotechnology Information (NCBI) Gene Expression Omnibus (GEO) under the respective accession numbers. Whole-cell lysate and secreted proteomics data are available at the PRIDE database under accession number PXD067052. Whole-exome sequencing data are available at the Sequence Read Archive (SRA) database under accession number SUB15543889.

## FUNDING INFORMATION

This paper was funded by CIHR project Grant (FRN 173360), a CRS operating grant, an Alberta Innovates Health Solutions Translational Research Chair and a Canada Research Chair awarded to LMP. MC was supported by a Canadian Institutes of Health Research (CIHR) Frederick Banting and Charles Best Canada Graduate Scholarship - Doctoral Award and a fellowship from Alberta Innovates Health Solutions.

---

**Supplementary Figure 1.**
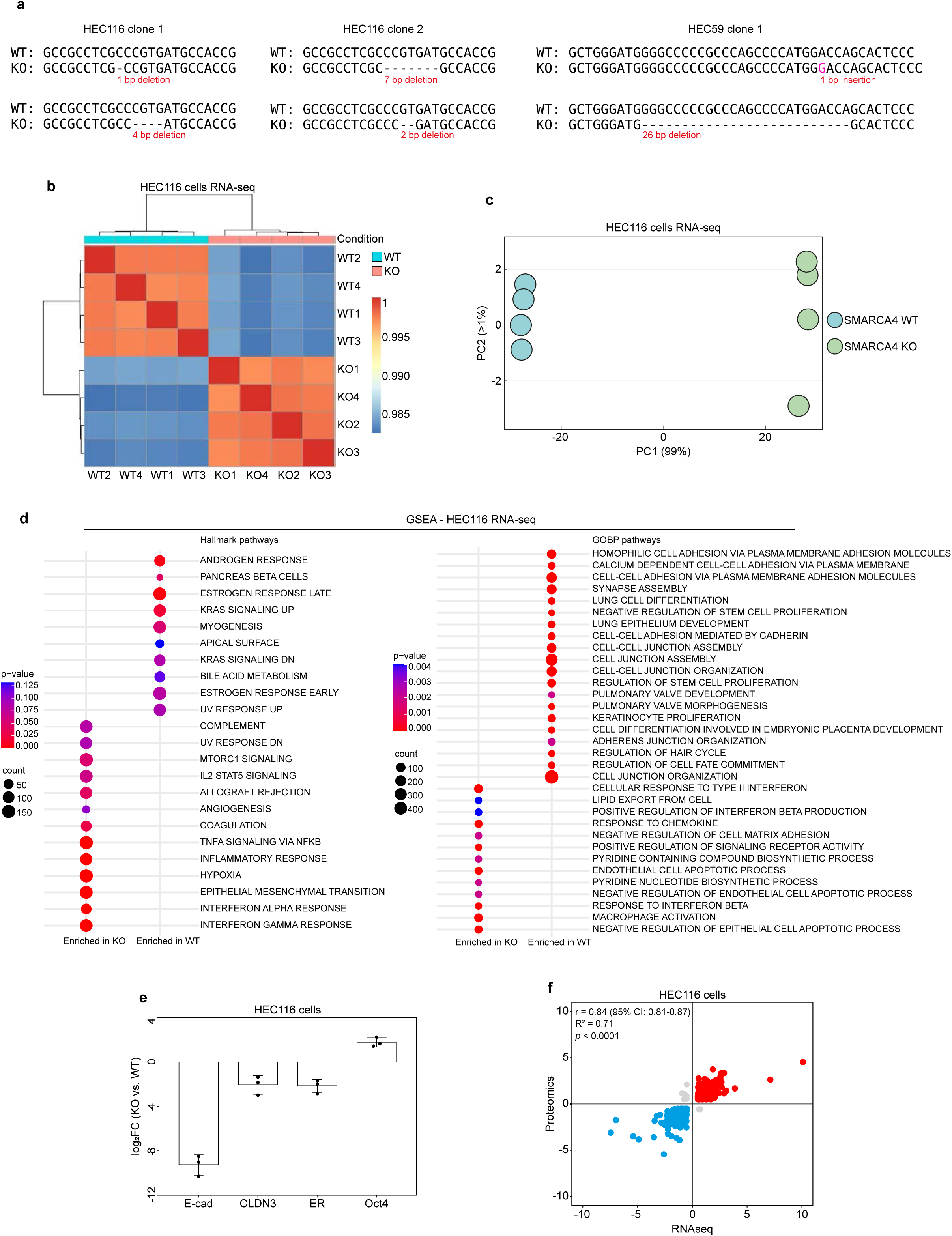
(a) CRISPR-Cas9-mediated SMARCA4 mutations in HEC116 and HEC59 clones. Shown are two HEC116 clones and one HEC59 clone, depicting the sequence of both alleles for each clone. The top row for each clone allele shows the WT reference sequence, and the bottom row shows the corresponding SMARCA4 KO sequence. Insertions and deletions (indels) introduced by CRISPR-Cas9 are highlighted. (b) Pairwise correlation heatmap of bulk RNA-seq profiles from HEC116 WT and SMARCA4 KO cells (n = 4). Colours indicate global expression similarity (orange = high, blue = low). Samples cluster according to global transcriptome similarities. (c) Principal component analysis (PCA) of transcriptomes upon SMARCA4 loss. PCA of bulk RNA-seq data from serum-starved HEC116 WT and SMARCA4 KO cells (n = 4). (d) Gene set enrichment analysis (GSEA) was performed on genes significantly regulated in HEC116 SMARCA4 KO versus WT cells (log₂FC > 1; *p* < 0.05). Top 20 pathways from Gene Ontology Biological Processes (GOBP) and all Hallmark pathways are shown. Dot size indicates the number of genes per pathway, and dot colour indicates FDR (all < 0.25). Enriched pathways in SMARCA4 KO versus WT cells are highlighted. (e) qPCR analysis of gene expression in HEC116 cells. Select markers of epithelial identity (*CDH1*/ E-CADHERIN), tight junctions (*CLDN3*), gynecological function (*ESR1* / ER), and stemness (*POU5F1* / OCT4) were quantified. Data represent log₂FC in SMARCA4 KO relative to WT controls from three independent replicates. (f) Correlation between transcriptome and proteome in HEC116 SMARCA4 KO cells. Scatter plot compares log₂FC from bulk RNA-seq and MS-based whole-cell proteomics (KO relative to WT). Pearson correlation indicates concordance between transcriptomic and proteomic changes (*p* < 0.001).

**Supplementary Figure 2.**
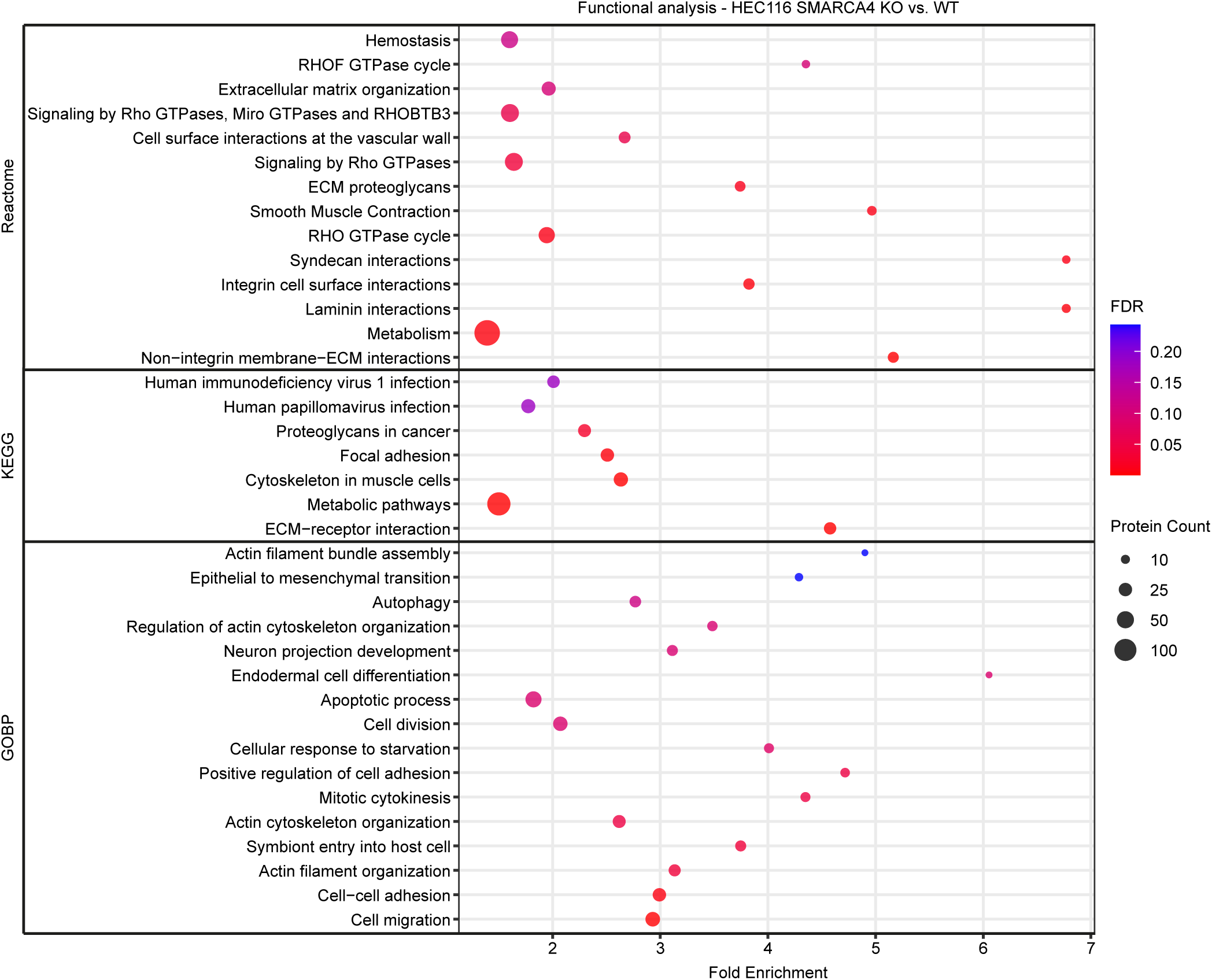
Functional enrichment of MS-based whole-cell proteome in HEC116 SMARCA4 KO cells versus WT. Over-representation analysis (ORA) was performed on significantly regulated proteins (log₂FC > 1; *p* < 0.05). Enriched pathways with FDR < 0.25 from REACTOME, Gene Ontology Biological Processes (GOBP), and KEGG are shown. Dot size indicates the number of proteins per pathway, and dot colour reflects the adjusted p-value.

**Supplementary Figure 3.**
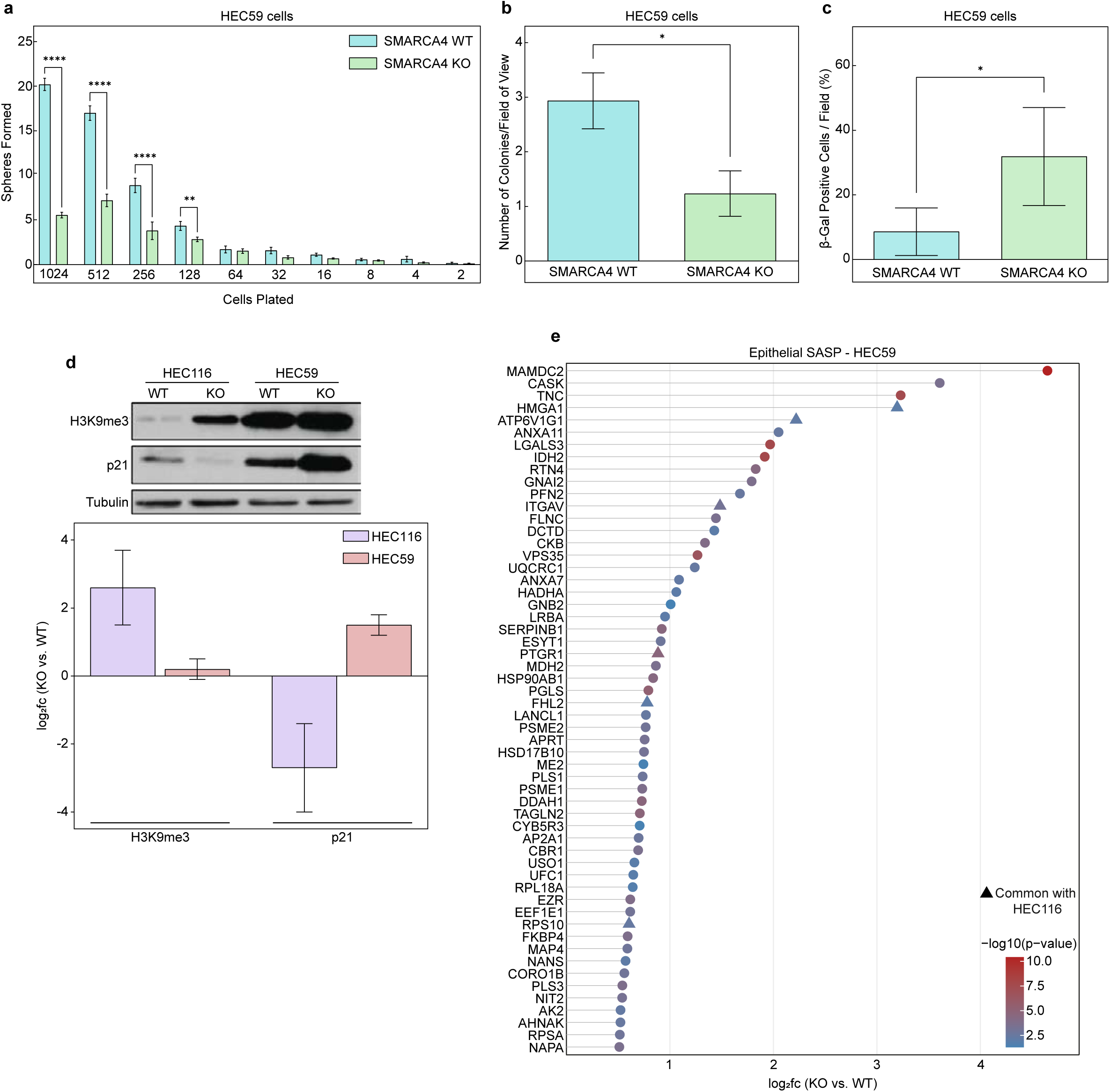
(a) Limiting dilution sphere formation assay. Single-cell suspensions of HEC59 WT and SMARCA4 KO were seeded at decreasing densities in ULA plates and cultured in minimal media for up to two weeks. Data represent eight technical replicates from three independent experiments. Multiple t-tests with Holm-Šidák correction assessed statistical significance (\*\**p* < 0.01, \*\*\*\**p* < 0.0001). (b) Soft agar colony formation assay. HEC59 WT and SMARCA4 KO cells were embedded in 0.7% agarose-containing media and cultured for up to two weeks. Colonies were counted from ten fields of view across three independent experiments. Statistical significance was assessed by unpaired t-test (\**p* < 0.05). (c) SA-β-gal assay. The percentage of positive cells in HEC59 WT and SMARCA4 KO cells was quantified from six randomly selected fields. Statistical significance was assessed by unpaired t-test (\**p* < 0.05). (d) Western blot analysis of H3K9me3 and p21 in HEC116 and HEC59 WT and SMARCA4 KO cells. Tubulin was used as a loading control. Representative blots from three independent experiments are shown. Densitometry analysis of the blots (bottom) was performed using ImageJ. Band intensities of H3K9me3 and p21 were normalized to tubulin, and log₂FC was calculated for KO relative to WT. Data represent the mean of three independent biological replicates. (e) MS-based secretome analysis. Proteins in CM of HEC59 SMARCA4 KO compared to WT with log₂FC > 0.5 and *p* < 0.05 were compared to the epithelial SASP signature (http://saspatlas.com). Colour scale indicates p-values of upregulated proteins. Proteins upregulated in both HEC59 and HEC116 SMARCA4 KO vs. WT are marked with ▴.

**Supplementary Figure 4.**
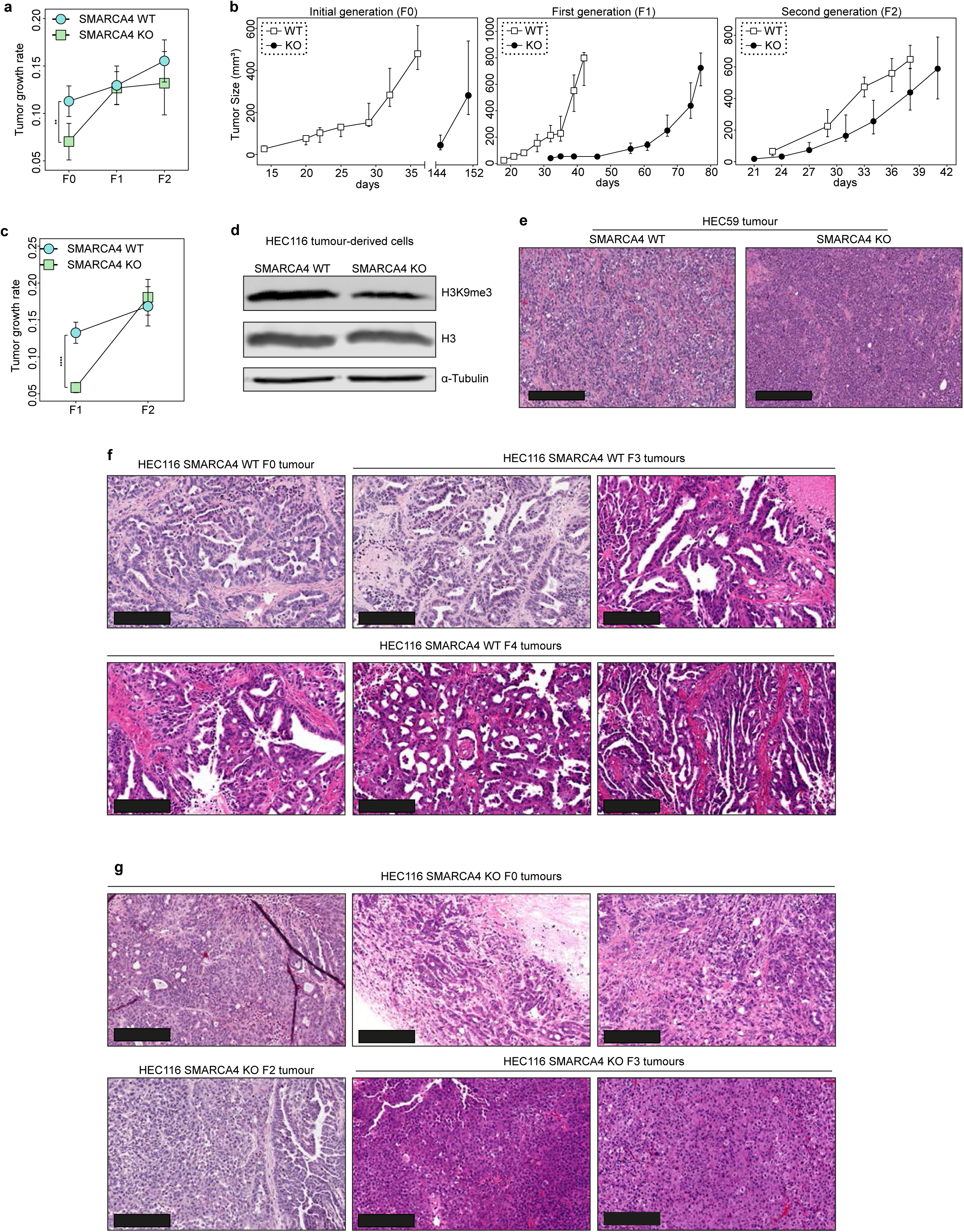
(a) Tumour growth rate (TGR) of serially passaged HEC116 (n = 6) WT and SMARCA4 KO CDX tumours across F0, F1, and F2 passages. TGR was calculated as the slope of a linear regression of the natural log of tumour volume over time, with higher slopes indicating faster growth. Statistical significance was assessed by multiple t-test with Holm–Šídák correction (\*\**p* < 0.01). (b) Tumour growth kinetics of HEC59 WT and SMARCA4 KO cells. Tumour cells were injected subcutaneously into NSG mice, and tumour area was measured over time across F0, F1, and F2 passages (n = 10 mice per group). (c) TGR of serially passaged HEC59 (n = 10) WT and SMARCA4 KO CDX tumours across F0, F1, and F2 passages. TGR was calculated as the slope of a linear regression of the natural log of tumour volume over time, with higher slopes indicating faster growth. Statistical significance was assessed by multiple t-test with Holm–Šídák correction (\*\*\*\**p* < 0.0001). (d) Western blot of H3K9me3 in tumour-derived HEC116 WT and SMARCA4 KO cells cultured *in vitro*. Representative blots from three independent experiments are shown. (e) Representative H&E images of second-generation (F2) HEC59 WT and SMARCA4 KO. Unlike HEC116 tumours, HEC59 tumours acquire an undifferentiated histology after serial passaging. Scale bar: 300 μm. (f) Representative H&E images of HEC116 SMARCA4 WT tumours across F0, F3 and F4 serial passages *in vivo* showing retention of endometrioid glandular-like histological features over time. Scale bar: 200 μm. (g) Representative H&E images of HEC116 SMARCA4 KO tumours across F0, F2 and F3 serial passages *in vivo* showing variable levels of dedifferentiation over time. Scale bar: 200 μm.

**Supplementary Figure 5.**
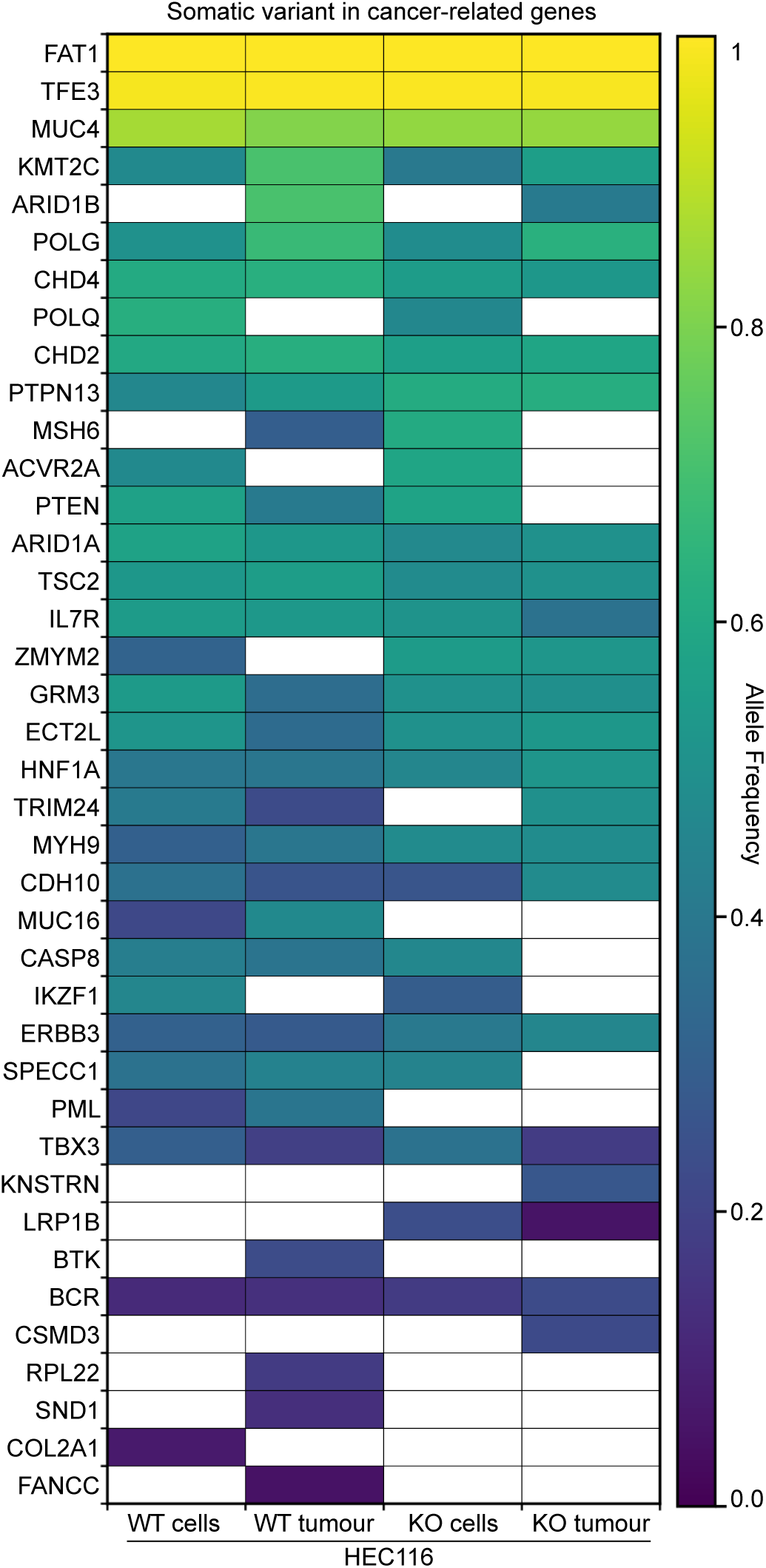
Heatmap of allele frequencies of nonsynonymous variants in cancer-related genes in HEC116 WT and SMARCA4 KO cells after three serial passages in mice. Variants are putatively somatic due to lack of paired normal samples. Genes are defined according to the COSMIC Cancer Gene Census. Colours indicate allele frequency, with white representing absence of a mutation.

**Supplementary Figure 6.**
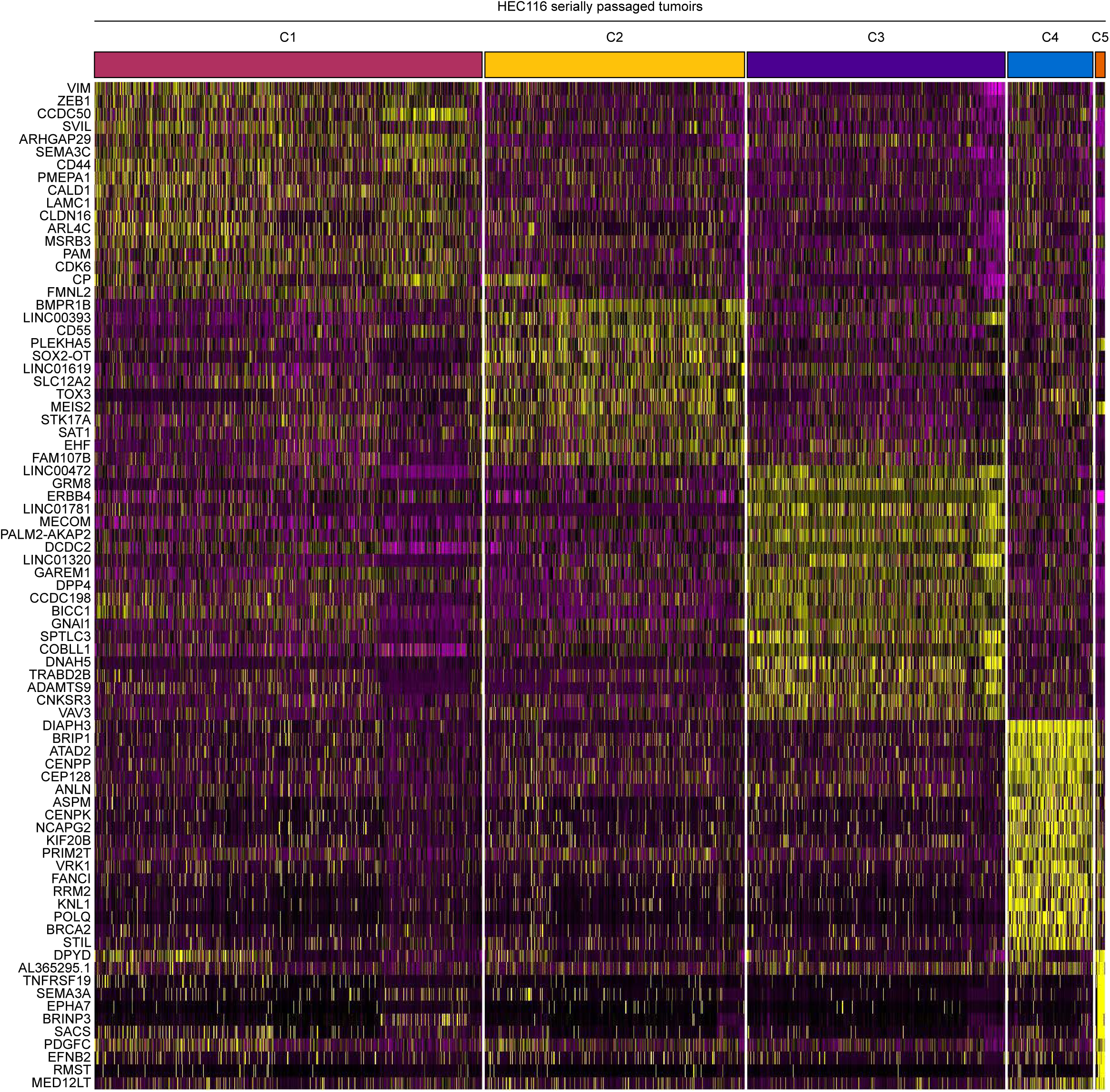
Heatmap of cluster-defining genes from Harmony-integrated scRNA-seq analysis of serially passaged HEC116 WT and SMARCA4 KO tumours. Rows represent transcriptional clusters (C1–C5) and columns show top genes in each cluster.

**Supplementary Figure 7.**
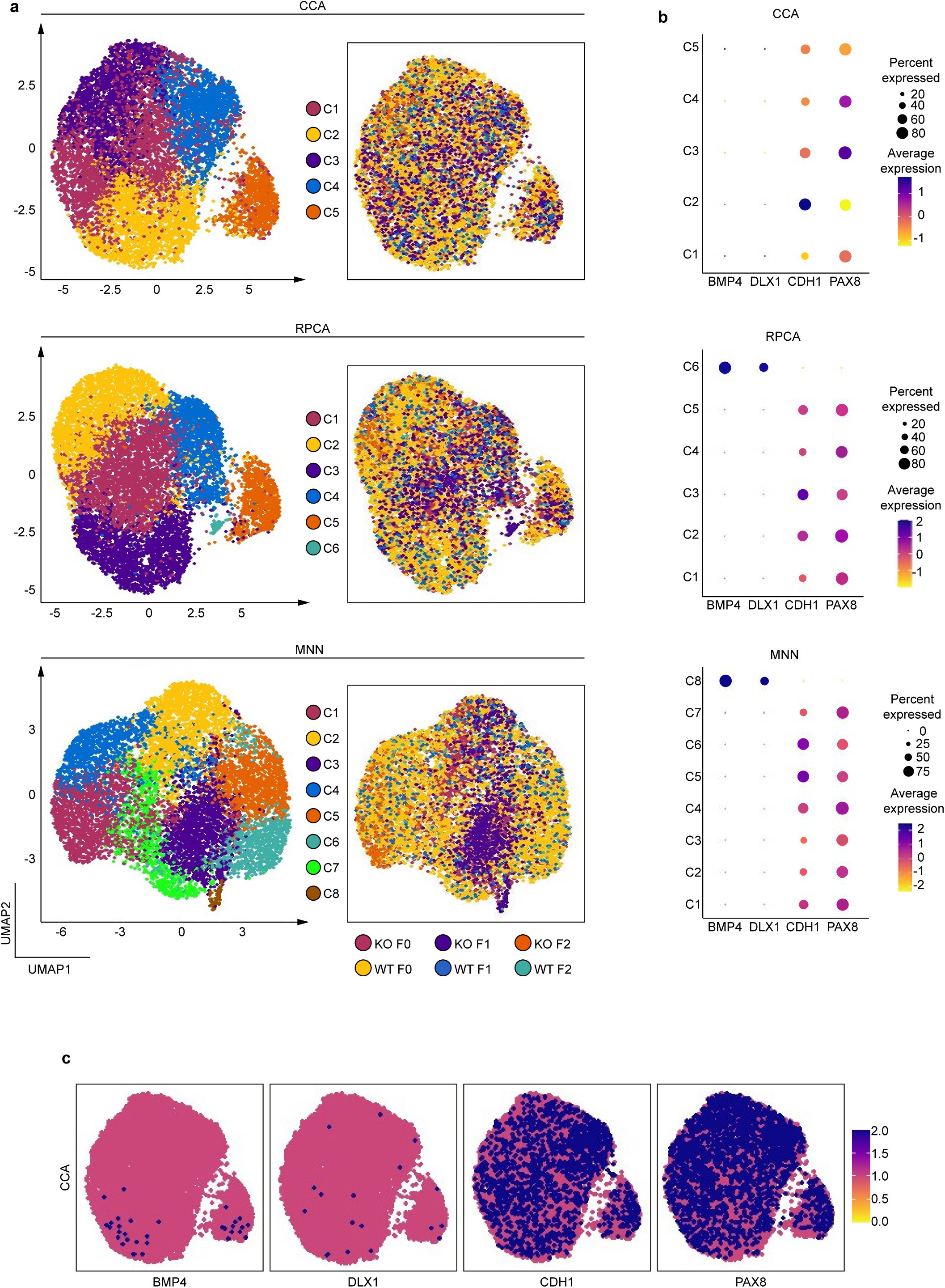
(a) Transcriptional clustering of serially passaged HEC116 WT and SMARCA4 KO tumours from scRNA-seq using alternative integration methods. Panels show cluster assignments for each integration method (CCA, RPCA, MNN) and sample composition within each cluster. (b) Dot plots of DDEC-associated marker expression across alternative integration methods from Harmony-integrated scRNA-seq analysis of HEC116 WT and SMARCA4 KO serially passaged tumours. Average expression of *BMP4*, *DLX1*, *CDH1*, and *PAX8* is shown for each cluster identified using CCA, RPCA, and MNN. Dot size indicates the percentage of cells in each cluster expressing the gene, and colour intensity indicates average expression. (c) Feature plots of DDEC-associated markers from scRNA-seq using CCA integration. Expression of *BMP4*, *DLX1*, *CDH1*, and *PAX8* is shown for individual cells from HEC116 WT and SMARCA4 KO tumours.

**Supplementary Figure 8.**
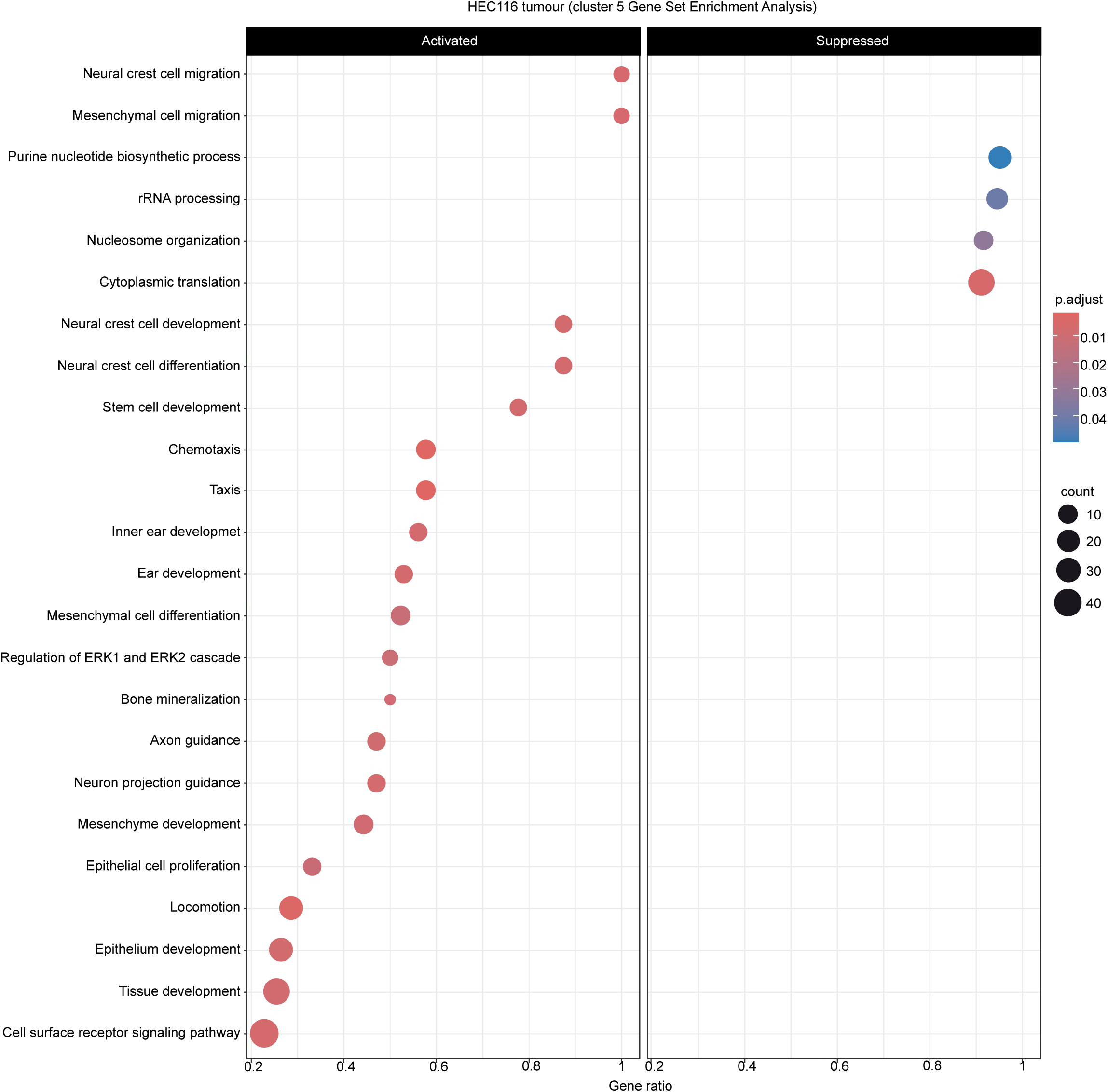
GSEA of the small cluster (C5) identified in HEC116 SMARCA4 KO tumours using Harmony integration. Each dot represents a gene set, with dot size showing the number of genes in the cluster and colour intensity indicating the p-value.

**Supplementary Figure 9.**
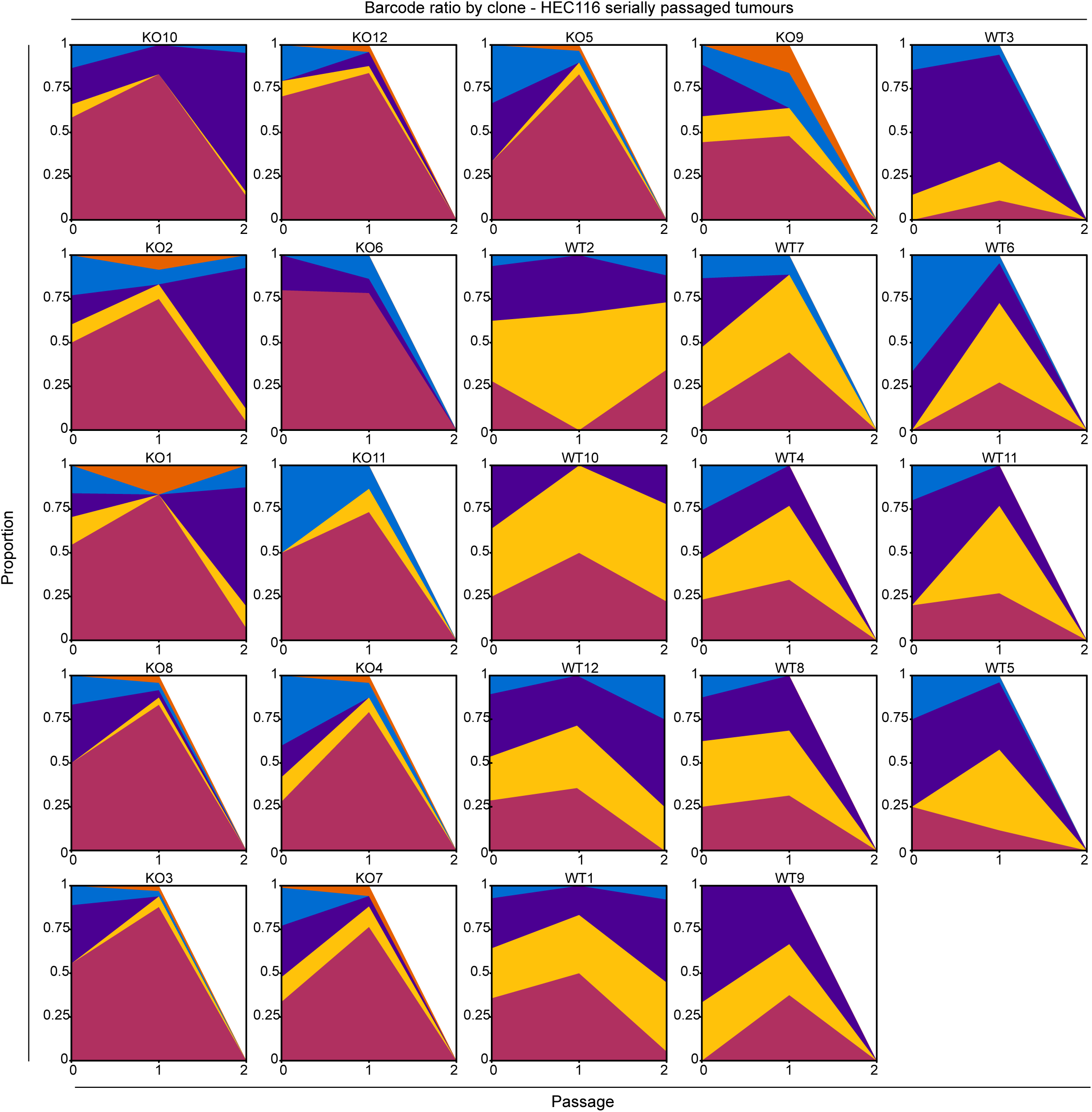
Clonal composition of HEC116 WT and SMARCA4 KO F0, F1, and F2 tumours. Individual clones were barcoded, and their distribution across transcriptome clusters (C1-C5) identified by Harmony-integrated scRNA-seq analysis is shown. Each line represents the proportional presence of a single clone across clusters over serial passages.

**Supplementary Figure 10.**
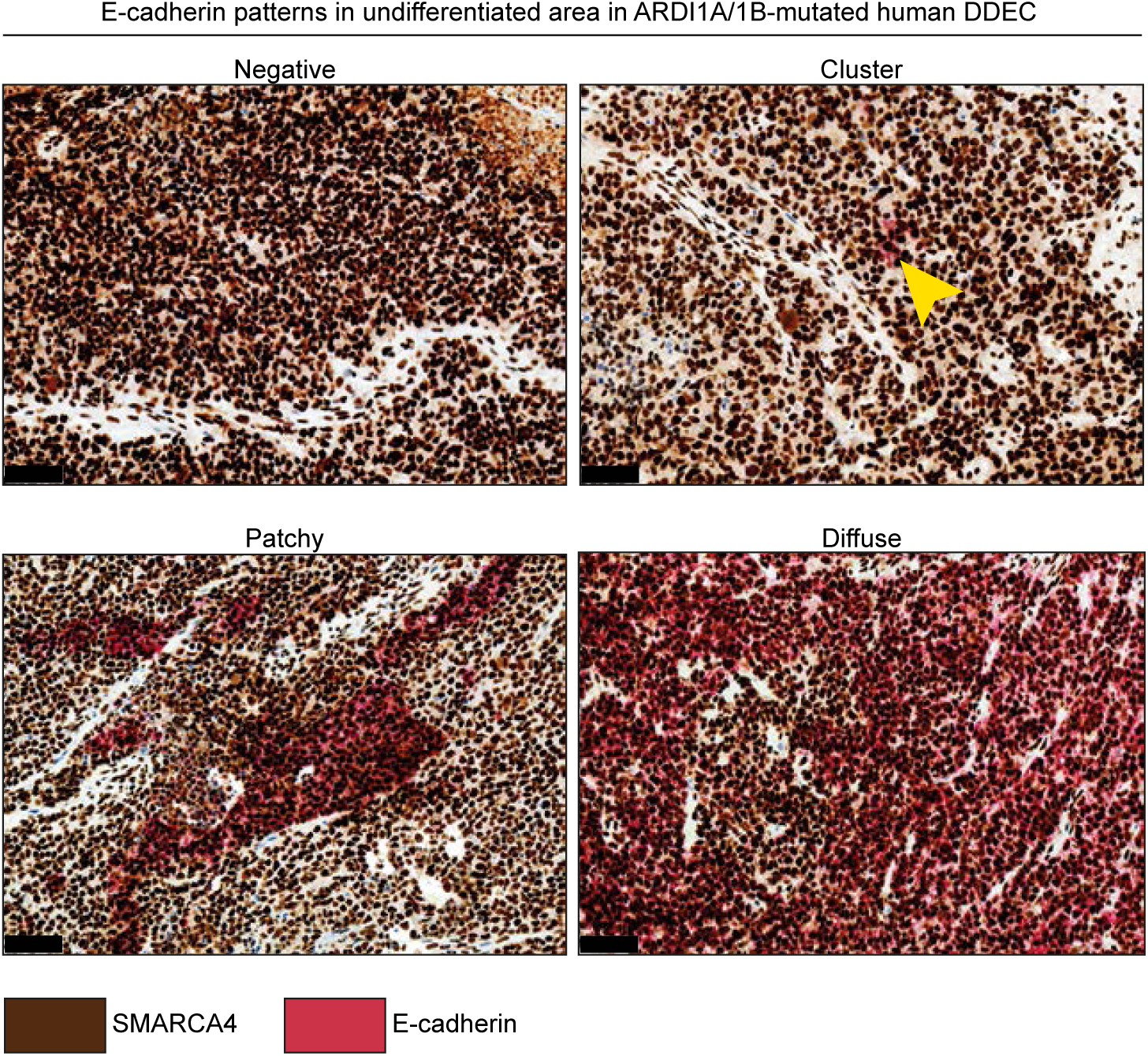
E-CADHERIN staining in poorly differentiated regions of human tumours with *ARID1A* or *ARID1B* mutations. Representative IHC images show distinct E-CADHERIN patterns: negative, clustered, patchy, and diffuse. Scale bar: 100 µm.

**Supplementary Figure 11.**
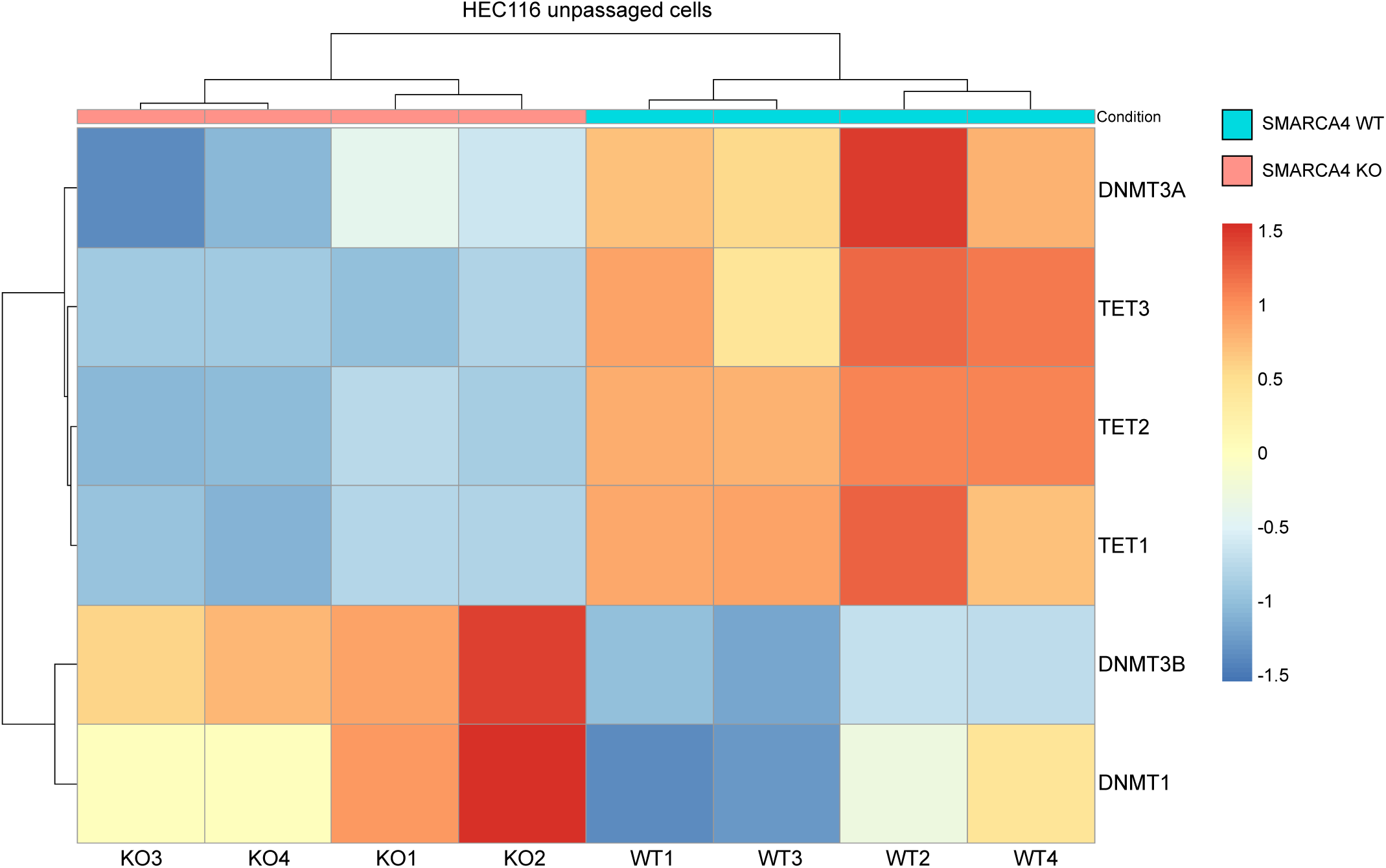
Heatmap showing expression of DNA methylases (*DNMT1*, *DNMT3A*, and *DNMT3B*) and demethylases (*TET1*, *TET2* and *TET3*) from bulk RNA-seq of HEC116 WT and SMARCA4 KO cells. Columns represent cell lines and rows represent genes.

## Notes

The authors declare no potential conflicts of interest.

### Competing Interest Statement

The authors have declared no competing interest.

